# One-Pot Isothermal Linear Amplification and Cas12a-based Nucleic Acid Detection

**DOI:** 10.1101/2025.06.27.661965

**Authors:** Selma Sinan, Remy M. Kooistra, Karunya Rajaraman, Zeba Islam, Damian Madan, Eric A. Nalefski, Ilya J. Finkelstein

**Affiliations:** Department of Molecular Biosciences and Institute for Cellular and Molecular Biology, University of Texas at Austin, Austin, Texas 78712, USA; Global Health Labs, Inc, Bellevue, WA 98007, USA; Institute for Protein Innovation, Boston, MA 02125, USA; Inari Agriculture Inc., Cambridge, MA 02139, USA; Center for Systems and Synthetic Biology, University of Texas at Austin, Austin, Texas 78712, US

## Abstract

CRISPR-based nucleic acid diagnostics are a promising class of point-of-care tools that could dramatically improve healthcare outcomes for millions worldwide. However, these diagnostics require nucleic acid pre-amplification, an additional step that complicates deployment to low resource settings. Here, we developed CATNAP (<u>Ca</u>s *<u>t</u>rans*-<u>n</u>uclease detection of <u>a</u>mplified <u>p</u>roducts), a method that integrates isothermal linear DNA amplification with Cas12a detection in a single reaction. CATNAP uses a nicking enzyme and DNA polymerase to continuously generate single-stranded DNA, activating Cas12a’s *trans*-cleavage activity without damaging the template. We optimized enzyme combinations, buffer conditions, and target selection to achieve high catalytic efficiency. CATNAP successfully distinguished between high- and low-risk HPV strains and detects HPV-16 in a cervical cancer crude cell lysate at room temperature with minimal equipment, offering advantages over PCR-based approaches. We conclude that CATNAP bridges the sensitivity gap in CRISPR diagnostics while maintaining simplicity, making accurate disease detection more accessible in resource-limited settings.

## Introduction

Rapid and accurate clinical tests are essential for effective patient care in low-resource settings where access to sophisticated diagnostic infrastructure is limited. CRISPR-based point-of-care tests that follow REASSURED criteria (real-time connectivity, equipment-free, affordable, sensitive, specific, user-friendly, rapid, environmentally friendly, deliverable) are emerging as powerful tools that can detect pathogen nucleic acids with high sensitivity in complex samples^1,2^. These assays leverage the *trans*-cleavage activity of Cas12a and related CRISPR nucleases, wherein target recognition triggers collateral cleavage of nearby single-stranded DNA molecules^1,3,4^. The binding of Cas12a to a target sequence adjacent to a protospacer-adjacent motif (PAM) activates the enzyme’s non-specific nuclease activity, resulting in the degradation of labeled reporter molecules and generation of detectable fluorescent signals^5^. This mechanism enables the development of highly specific and sensitive detection platforms for pathogen identification in low-resource settings^6,7^.

Most pathogens exist at attomolar to femtomolar concentrations in patient specimens, but Cas12a-based diagnostics^8,9^ are typically sensitive to nucleic acids that are in the picomolar to nanomolar range^1,10^. This sensitivity gap necessitates pre-amplification of target nucleic acids prior to CRISPR-based detection. Common pre-amplification strategies include Polymerase Chain Reaction (PCR)-based methods and isothermal techniques such as Loop-Mediated Isothermal Amplification (LAMP) or Recombinase Polymerase Amplification (RPA). However, these approaches present significant challenges in low-resource settings^11,12^. Specialized equipment, trained personnel, and reliable power sources are often unavailable. Reagent costs and cold-chain requirements further restrict accessibility. PCR requires thermal cycling, restricting its utility for point-of-care applications. Isothermal methods also require complex primer design, suffer from non-specific amplification, and are challenging to convert into a quantitative assay^13^. These protocols increase complexity and time demands, thus reducing the feasibility of rapid testing. Further, contamination risks during exponential amplification are particularly problematic in settings with limited laboratory infrastructure.

Here, we introduce CATNAP (Cas *trans*-nuclease detection of amplified products), a one-pot diagnostic assay that combines isothermal linear DNA amplification with Cas12a-based detection. Linear amplification reduces contamination risk, has higher specificity, and is simpler to quantify relative to exponential amplification methods. CATNAP uses a site-specific nicking endonuclease and strand-displacing DNA polymerase to continuously generate single-stranded DNA from double-stranded templates. These ssDNAs activate Cas12a’s *trans*-cleavage activity^14^ without damaging the original template DNA, thereby resulting in continuous signal amplification. By strategically designing crRNAs that target regions with dipurine PAMs near nicking sites, we achieved rapid and specific detection of HPV-16 DNA with high catalytic efficiency. We demonstrated CATNAP’s ability to discriminate between oncogenic and low-risk HPV strains and successfully detected HPV-16 in cervical cancer cell lysates. The method’s compatibility with room temperature operation and simplified workflow makes it particularly suitable for point-of-care diagnostics in resource-limited settings where sophisticated equipment and technical expertise are unavailable. More broadly, combining linear pre-amplification with sensitive detection will accelerate the development of point-of-care diagnostics in resource-limited settings.

## Results

### Development of a Cas12a-Based One-Pot Amplification and Detection Assay

**Figure 1** illustrates CATNAP (<u>Ca</u>s *<u>t</u>rans*-<u>n</u>uclease detection of <u>a</u>mplified <u>p</u>roducts). CATNAP integrates isothermal linear amplification with CRISPR RNA (crRNA)-guided Cas12a ribonucleoprotein (RNP)-based detection. Targeted nucleic acid amplification relies on the coordinated activities of a site-specific nicking endonuclease (nickase) and a strand-displacing DNA polymerase (DNAP). First, the nickase cuts the target DNA strand downstream of the Cas12a target. A DNA polymerase with strand displacement activity then extends this nick, synthesizing a new DNA strand while displacing the downstream target single-stranded DNA (ssDNA). Polymerase activity also re-generates the nick site for subsequent re-nicking and re-synthesis. The crRNA guides Cas12a to this ssDNA target, activating Cas12a’s *trans*-cleavage activity^15^. Once activated, Cas12a cleaves fluorescently labeled *trans*-substrates in the reaction mixture, producing a detectable signal. We designed protospacers containing central dipurines in their non-target strand (NTS) PAMs because Cas12a can be activated on these targets, but only when they are ssDNA (see below)^16^. Multiple rounds of concurrent nicking and strand displacement linearly amplify the ssDNA, enabling detection of low-abundance DNA sequences with high specificity and sensitivity.

**Figure 1.**
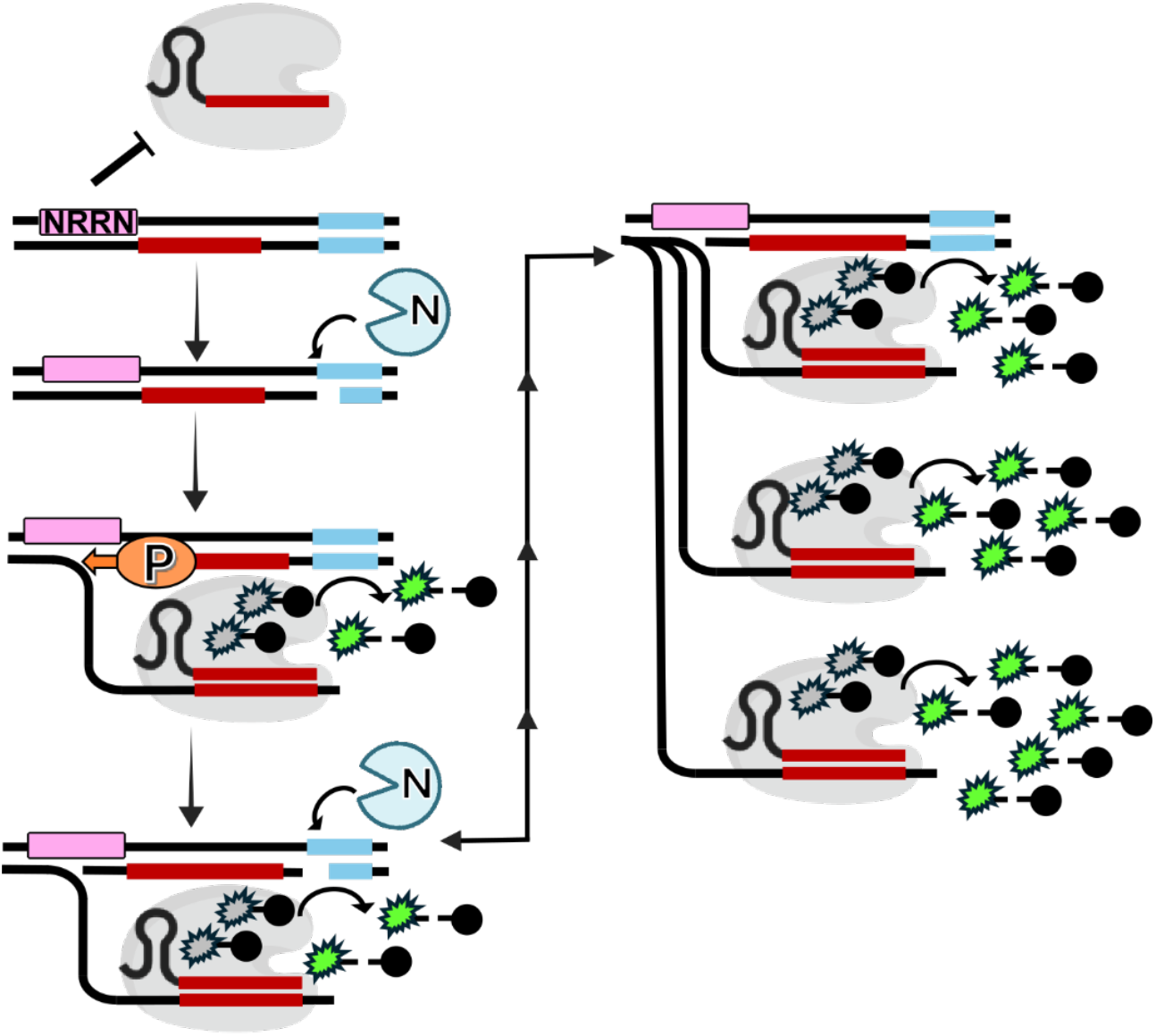
Schematic of linear DNA amplification and detection. A Cas12a target (red) is downstream of a nicking site (blue) and is flanked by a non-ideal PAM containing central dipurines (NRRN, pink). This PAM prevents Cas12a binding to the double-stranded DNA. A nicking enzyme (N, blue) creates a nick upstream of the target site (red) on the bottom strand, generating a priming site. A DNA polymerase (orange, P) extends the nick via strand displacement synthesis, producing single-stranded DNA (ssDNA). This ssDNA activates the crRNA-Cas12a ribonuclear protein (RNP, gray), inducing the RNP to cleave the reporter substrate in *trans* (green-black). Further DNA polymerase activity restores the nicking site, allowing the cycle to repeat and amplify the reporter signal.

### Benchmarking CATNAP for Detecting Human Papillomavirus

We first tested whether Cas12a RNPs could cleave plasmid DNA at HPV-16 target sites containing either dipurine or dipyrimidine PAMs (**Table S1**). We targeted conserved viral genes in HPV-16, because this strain is strongly linked with cervical cancer and is an important target for developing a fieldable diagnostic^17,18^. Because CATNAP continuously resynthesizes the target strand, we also selected targets that were upstream of a Nt.BsmAI nickase site (**Table S1**).

We incubated the RNPs with plasmids encoding HPV targets with different PAMs at 37°C. RNPs targeting dipyrimidine PAMs cleaved the plasmid and produced a linear DNA band (**Figure 2A**). In contrast, RNPs targeting dipurine PAMs (AGAT and GGGT) did not cleave the plasmid, even after 30 minutes. This result confirms that dipurine PAMs do not support Cas12a activation on plasmid DNA. We previously reported RNPs targeting dipurine PAMs are efficiently activated by short ssDNA but not dsDNA targets^16^ and used this insight to prioritize HPV target sites that selectively trigger Cas12a activity as ssDNA but not in double-stranded form. As expected, RNPs targeting dipurine PAMs were activated by ssDNA, but not dsDNA (**Figure 2B, S1A and Table S2**)^16^. In contrast RNPs targeting dipyrimidine PAMs were activated by both dsDNA and ssDNA (**Figures S1A, B**).

**Figure 2.**
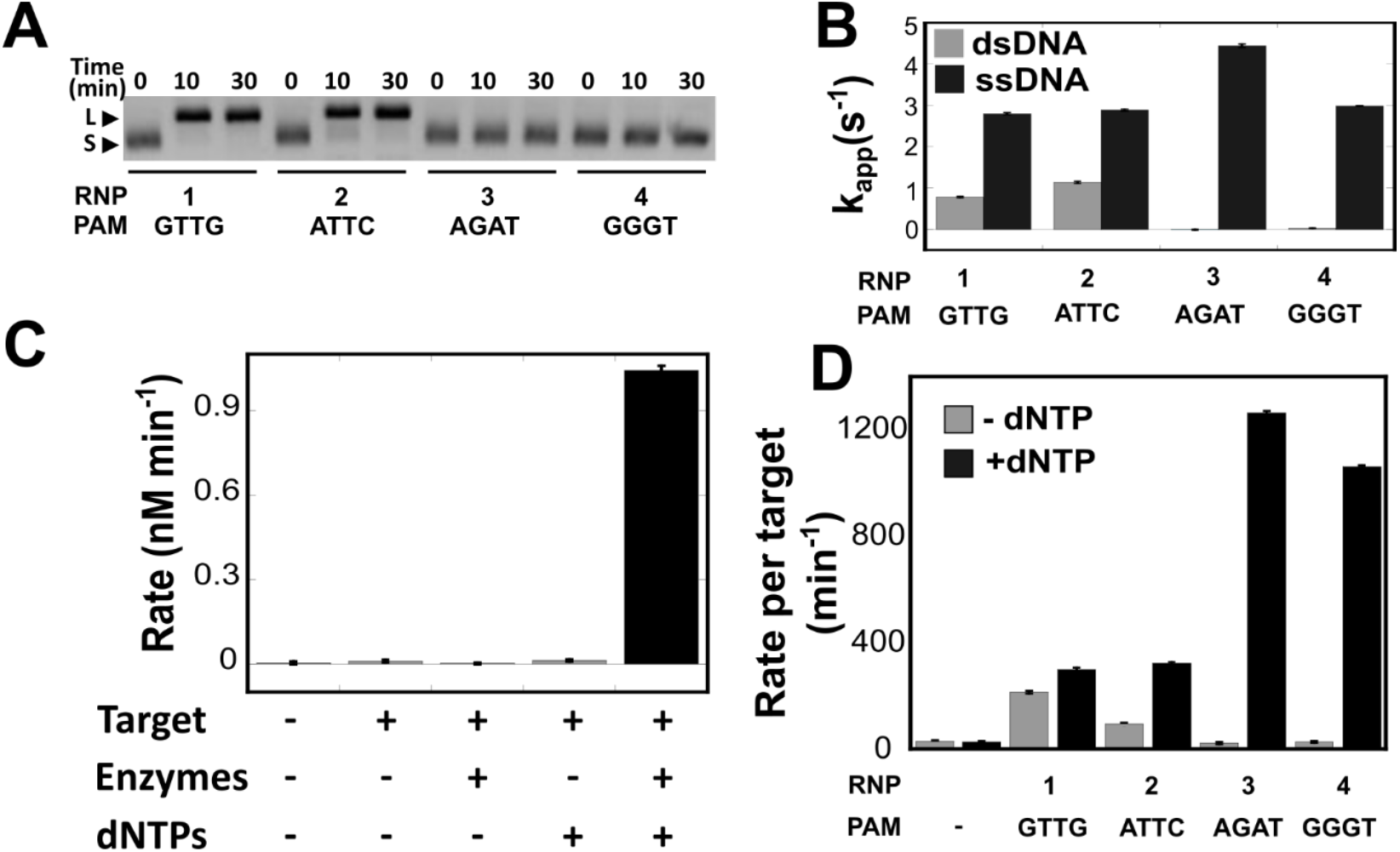
CATNAP validation and assay requirements. (A) Cas12a efficiently cuts targets with dipyrimidine, but not dipurine PAMs. RNPs were incubated with plasmid DNA containing target sequences, and at the indicated times, reactions were quenched and subjected to electrophoresis on a 1.5% agarose gel containing ethidium bromide followed by visualization. Linear (L) and uncleaved supercoiled (S) DNA bands are indicated by arrowheads. (B) RNPs using dipurine PAMs are efficiently activated by ssDNA but not by dsDNA targets. Bars (± SE) represent apparent turnover (k_app_) of *trans*-substrate FQ-C_10_ for HPV-16-specific RNPs reacted in triplicate with protospacer targets within long, 858-bp dsDNA (Target-E of HPV-16) or short, 40-nt ssDNA (see **Figure S1 and Table S2**). (C) CATNAP signals are generated with the correct combination of target DNA, reaction enzymes and dNTPs. Cleavage of FQ-C_10_ in one-pot reactions by an RNP with the indicated CATNAP components, where bars represent mean (± SE) *trans*-cleavage rates for triplicates (see **Figure S2 and Table S**4). (D) CATNAP activates *trans*-substrate cleavage by RNPs utilizing dipurine PAMs. CATNAP reactions were performed in the presence of a plasmid encoding a complete HPV-16 genome. Bars represent reporter cleavage rates (± SE) for triplicates (see **Table S3**) normalized to the concentration of input target DNA. Signals generated from RNPs utilizing dipyrimidine PAMs in reactions lacking dNTPs reflect the expected direct activation by input target DNA.

Next, we implemented the CATNAP reaction using Nt.BsmAI nicking enzyme and the *E. coli* Klenow Fragment (exo-) of DNA polymerase I (DNAP; **Figures 2C-D, Figure S2, Tables S3**,**4**). We selected *E. coli* Klenow Fragment (exo-), a variant of DNA Polymerase I lacking 3’-5’ exonuclease activity, because it possesses moderate strand displacement capability^19^. Quantitative PCR confirmed efficient production of at least six ssDNA copies per template within 30 minutes in two different reaction buffers (**Figures S3**). As expected, efficient Cas12a activation required all enzymes and dNTPs to amplify the target ssDNA. Both the established *trans*-cleavage buffer^16^ and rCutSmart buffer supported effective ssDNA synthesis (**Figure S3B**). The rCutSmart buffer was specifically formulated to accommodate the enzymatic activities of both the nicking enzyme and polymerase, making it particularly suitable for CATNAP. Both buffers proved to be more effective than NEBuffer2, which is recommended for use with DNAP, in promoting Cas12a *trans*-substrate cleavage (**Figure S4**).

We next tested the relationship between RNP activity and distance separating the targeting sequences from the nicking site. Two-step reactions were performed using a panel of RNPs targeting non-overlapping sequences adjacent to a common nicking site in HPV-16 (**Table S5**). The first step consisted of ssDNA generation by nicking-polymerase reactions without added RNP, and in the second, *trans*-cleavage by the RNPs in response to the reaction products was measured (**Figure S5**). Rates of *trans*-cleavage correlated directly with proximity of the Cas12a targeting sequence to the nicking site, suggesting the efficiency of DNA synthesis decreases as distance from the nicking site increases, consistent with the limited strand displacement activity of Klenow DNAP.

We reasoned that the rates of both Cas12a *trans* cleavage and DNAP synthesis are the two critical parameters for efficient CATNAP amplification and detection. Cas12a orthologs exhibit significant diversity in their collateral DNase activities upon target recognition^20,21^. Therefore, we tested six additional Cas12a orthologs in the full CATNAP reaction (**Figures 3A and S6**)^20^. These orthologs encompass five enzymes from mammalian microbiota—*Butyrivibrio sp*. NC3005 (BsCas12a), *Bacteroidetes oral* taxon 274 (BoCas12a), *Eubacterium rectale* (ErCas12a), *Francisella novicida* U112 (FnCas12), and *Acidaminococcus sp*. BV3L6 (AsCas12a)—as well as *Thiomicrospira sp*. XS5 (TsCas12a), isolated from a brine-sea water interface. We tested each ortholog in the full CATNAP reaction with the FQ-C_10_ *trans*-substrate (**Figure 3A**). CATNAP *trans*-cleavage rates varied over a ~20-fold range, with BsCas12a showing the highest rate (54 ± 1 min^−1^), 1.4-fold higher than LbCas12a, the next best (**Figure S6**). The superior performance of BsCas12a in the CATNAP assay likely stems from an optimal balance between its target DNA binding affinity, *trans*-cleavage activation kinetics, and compatibility with the DNAP in the reaction buffer. To find the best RNP concentration for CATNAP, we ran reactions with different RNP amounts and crRNAs at a limiting level (**Figure S7**). We found that RNP concentrations above 8 nM caused inhibition of CATNAP detection.

**Figure 3.**
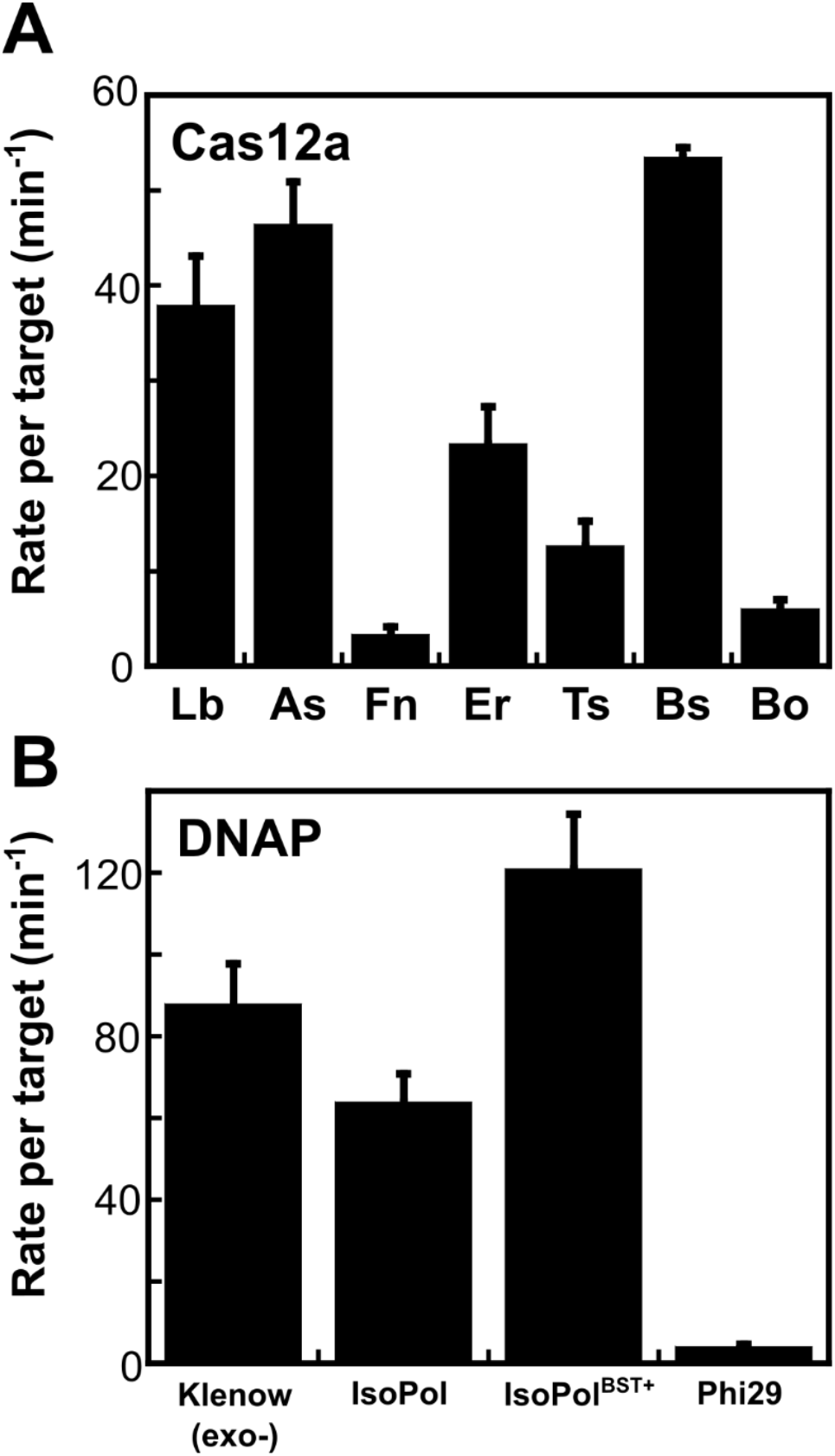
CATNAP component optimization. CATNAP reactions were carried out with RNPs assembled from different orthologs of Cas12a (A) or with different DNA polymerases (B). Symbols represent reporter cleavage rates (± SE) for triplicates normalized to the concentration of input target DNA from time courses in Figure S6 and Figure S8. All reactions were conducted with RNP-(3) and HPV-16 plasmid at 37°C.

We compared the performance of Klenow (exo-) against three other DNAPs in the full CATNAP reactions (**Figures 3B and S8**,**9**). All reactions included an Cas12a RNP targeting the HPV-16 genome (**Table S1**). IsoPol and IsoPol^BST+^ are two engineered DNAPs that exhibit excellent isothermal strand displacement activity in high ionic strength buffers, with IsoPol^BST+^ more heat-resistant above 37°C. We also tested phi29 DNAP for its excellent strand-displacement activity at 30°C. Klenow (exo-) and IsoPol DNAPs performed similarly within the full CATNAP reaction (**Figure 3B**). IsoPol^BST+^ showed the highest activity at 37°C, consistent with its mesophilic characteristics. Surprisingly, Phi29 DNAP completely inhibited the CATNAP reaction, likely due to competition with Cas12a for the generated ssDNA or its 3’-5’ exonuclease activity^22^. Based on these results and additional experiments that optimized both Cas12a and DNAP concentration, all further CATNAP experiments were conducted with 2.0 nM RNP and 62 U /mL Klenow fragment (exo-) DNAP.

### Discriminating Between Oncogenic and Low-Risk HPV Types

We evaluated whether CATNAP could distinguish closely related HPV types (**Figure 4**), including three high-risk types associated with cervical cancer (HPV-16, -18, and -52) and one low-risk type (HPV-6b). To achieve this, we identified distinct Nt.BsmAI restriction sites in the genomes of each type and designed crRNAs targeting dipurine PAM-associated protospacers near the nicking sites. As expected for CATNAP RNPs, each crRNA-guided complex demonstrated strong *trans*-cleavage activity when exposed to ssDNA sequences containing the correct protospacers, while showing no significant activity against plasmids encoding the respective full HPV genomes (**Figure S10)**. In one-pot CATNAP reactions, each RNP demonstrated high *trans*-cleavage activity specifically in response to its cognate HPV plasmid, confirming accurate viral genotyping (**Figure 4, Table S6-8**). We conclude that CATNAP can distinguish between highly-related pathogenic DNA targets.

**Figure 4.**
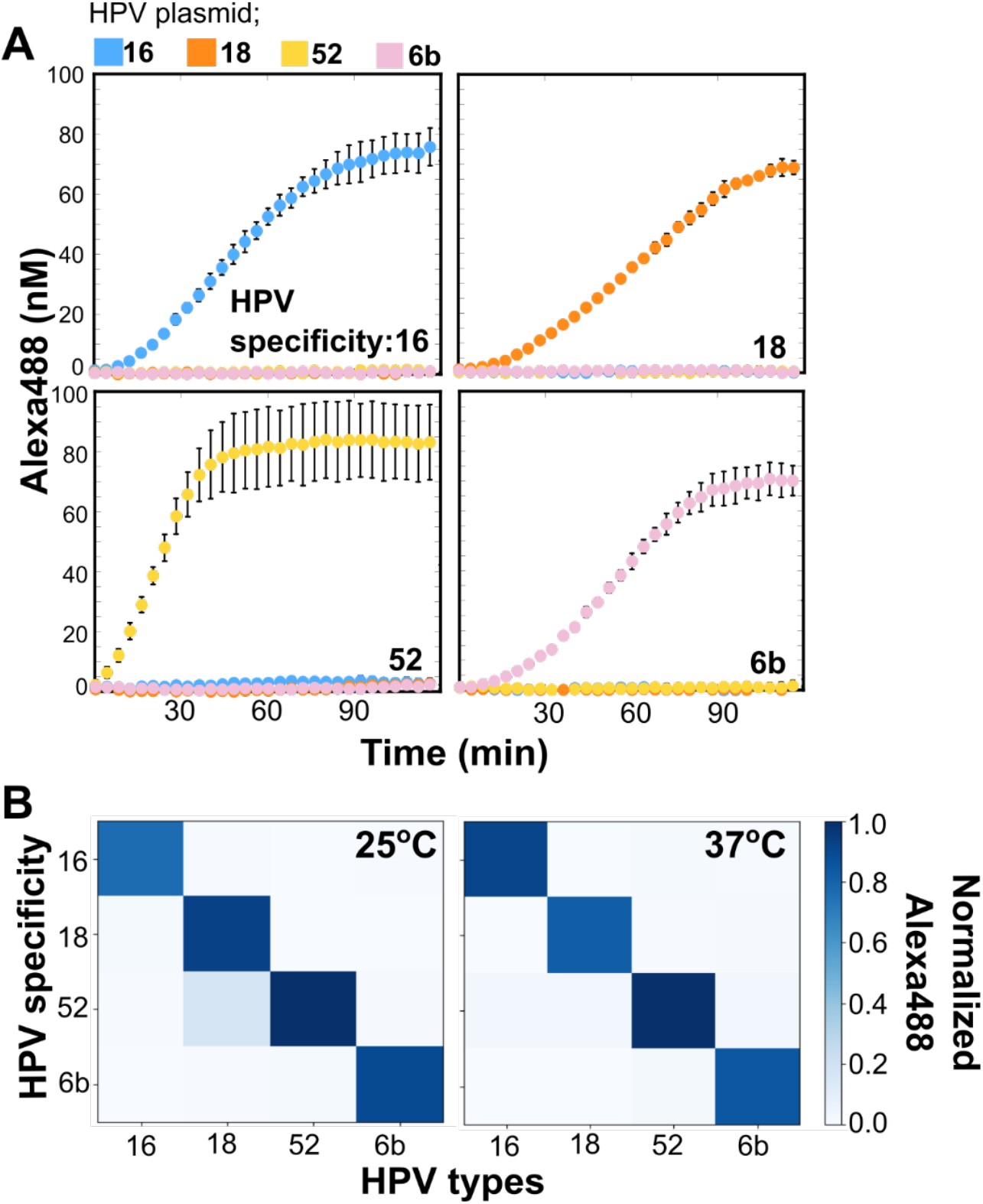
Type-specific HPV detection by CATNAP. (A) CATNAP reactions were performed at 37°C with RNPs designed to HPV-16, -18, -52, and -6b (**Figure S10**) using plasmids containing each type. Symbols represent mean (± SE) of background-corrected product measured in triplicate. (B) Normalized product accumulated at 2hr from CATNAP reactions, represented as a heat-map for 25°C and 37°C (**Table S6**,**7**), demonstrating specificity of each RNP (y-axis) for its intended target (x-axis).

Additionally, we observed that the specificity of these reactions was retained at 25°C (**Figure 4**), although the overall assay signal was reduced (**Figure S11**). Notably, BsCas12a, the top performer at 37°C, remained the most active ortholog at 25°C, yielding the highest *trans*-cleavage rate under our CATNAP conditions (**Figure S12)**. The preservation of activity across this temperature range suggests that CATNAP can be deployed in settings without the need for external heating equipment, thereby extending its potential utility for strain-specific viral detection in low-resource settings.

### Measuring analytical sensitivity of CATNAP

We measured the analytical sensitivity of CATNAP for detection of HPV-16 and compared it to amplification-free detection (**Figure 5**). Two different polymerases (IsoPol^BST+^ and Klenow exo-) and two Cas12a variants (BsCas12a or LbCas12a) were tested in CATNAP, combined with pools of 12 HPV-16-specific RNPs that utilize dipurine PAMs. For amplification-free detection, a pool of 20 HPV-16-specific RNPs that utilize dipyrimidine PAMs were used. Fluorescence values were collected over 2 hours after initiation of reactions. LODs were calculated at each time point using a logistic fit of the raw fluorescence data (**Figures 5A, S13**). With the same polymerase, BsCas12a was much faster than LbCas12a. Its slope was ~21 RFU m^−1^ fM^−1^, seven-fold higher than LbCas12a (3 RFU min^−1^ fM^−1^). The BsCas12a signal also plateaued at ~20 000 RFU instead of ~7,000 RFU, indicating a ~3-fold improvement. Keeping LbCas12a constant and swapping the polymerase showed the same pattern. IsoPol BST+ gave a slope of ~5 RFU min^−1^ fM^−1^, and Klenow exo-reached only ~3 RFU min^−1^ fM^−1^ (**Figures 5B, S14**). Figure of merit^23^ (FOM) was calculated from each LOD (**Figure 5B**). The results indicate that by 30–60 minutes, each of the three reactions had fully developed. At 60 minutes, the measured LODs were 0.57 ± 0.07 fM and 0.93 ± 0.16 fM for the IsoPol and Klenow CATNAP reactions, representing 3–5 times greater sensitivity than the 2.7 ± 1.1 fM for the amplification-free 20-RNP pool. Moreover, when CATNAP assays were performed at 25°C using both IsoPol BST and Klenow exo-, we observed similar LODs and reaction kinetics to those at 37°C, demonstrating that analytical sensitivity is maintained under reduced-temperature conditions (**Figure S15**).

**Figure 5.**
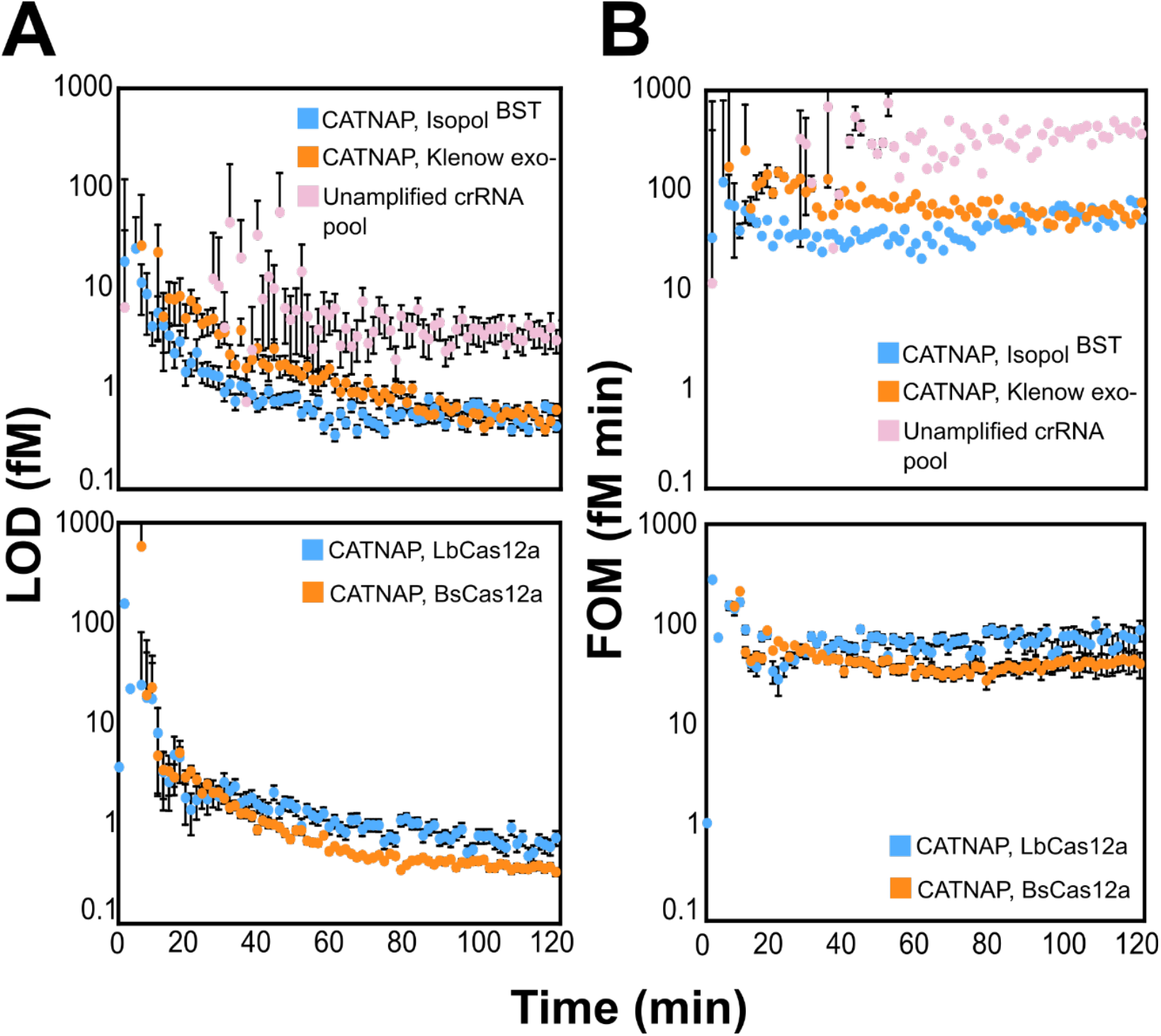
Limit of detection (LOD) and figure of merit (FOM). (A) CATNAP shows greater sensitivity for target detection than amplification-free detection by RNP pools. *Trans*-substrate reporter cleavage was measured in response to a titration of HPV-16 plasmid by either CATNAP employing two different DNA polymerases and pools of 12 CATNAP RNPs or by amplification-free detection using a pool of 20 non-CATNAP RNPs (top) or CATNAP using pools of 12 CATNAP RNPs formed from two different Cas12a orthologs (bottom). Symbols and bars represent mean (± SE) of LODs calculated at each time point based on the fluorescent signal (see **Figures S13, 14**). (B) Figure of Merit (FOM) comparison for HPV-16 plasmid detection by CATNAP versus amplification-free RNP pools. Symbols and bars represent the mean (± SE) of FOM values determined from LOD at each time point in A.

### HPV Detection in Cervical Cancer Cells

To evaluate CATNAP’s ability to detect HPV in a biologically relevant context, we tested lysates from two cervical cancer cell lines. CaSki (ATCC CRM-CRL-1550) contains integrated HPV-16 DNA, while C-33 A (ATCC HTB-31) is HPV-negative. Two crRNAs were used in CATNAP: one specific for HPV-16 (RNP-16 in **Table S1**) and a non-targeting control specific for *Mycobacterium ulcerans* (RNP-54 in **Table S1**). Reactions were with and without dNTPs to assess the impact of polymerase-driven amplification. As expected, RNP-473 was activated in the presence of Casksi lysates, but only with pre-amplification (**Figure 6**). Both CaSki and C33A lysates without amplification matched the responses of RNP-54, regardless of amplification (**Figures S16-S18 Table S9**). This indicates the background is a non-specific response of the RNP to genomic DNA. Despite the non-specific interactions with genomic DNA, CATNAP shows ~4-fold discrimination between HPV-positive and HPV-negative cells.

**Figure 6.**
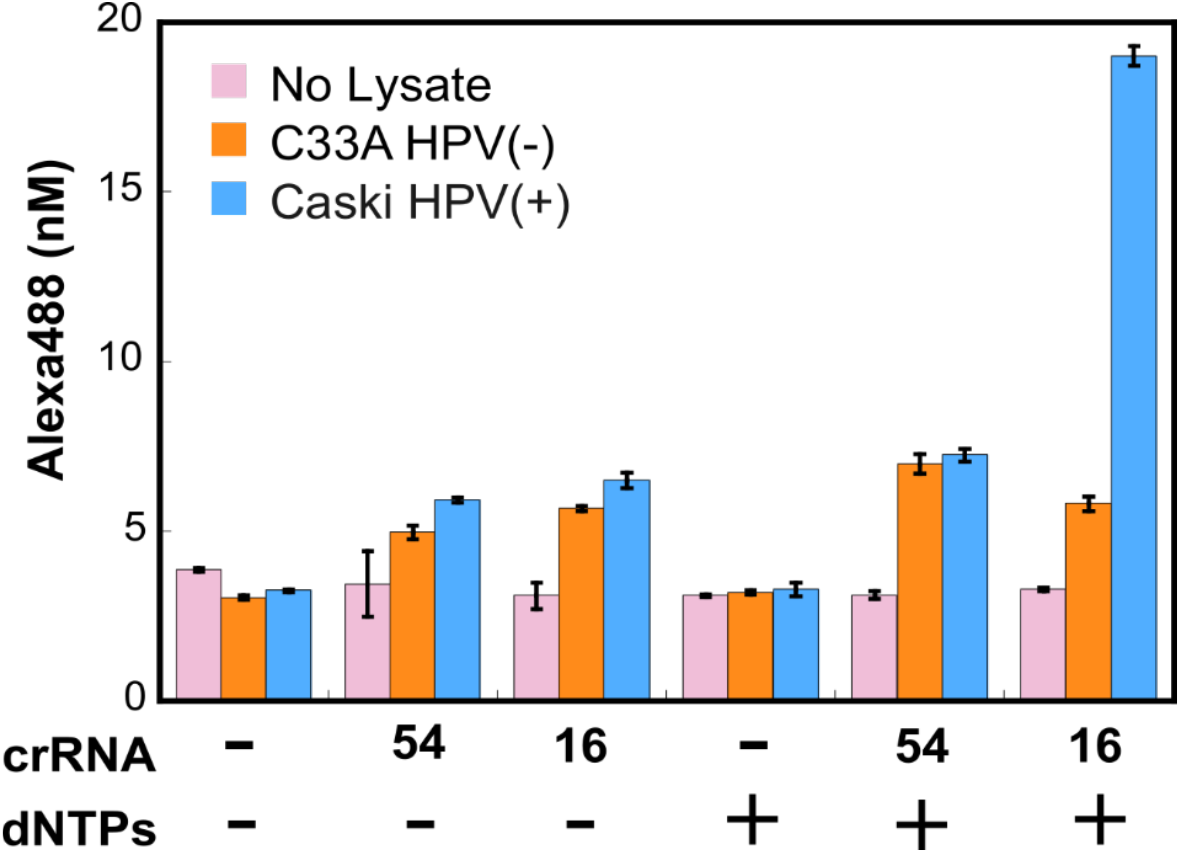
CATNAP-based HPV detection in crude cell lysates. CATNAP was performed on crude lysates prepared from C-33A or CaSki cells using an HPV-16-specific CATNAP RNP (RNP-16) and control conditions (no RNP or non-CATNAP RNP-54 specific to *M. ulcerans*). CaSki cervical cancer cells harbor HPV-16, whereas C-33A cervical cancer cells do not. Bars represent the mean (± SD) of product from triplicates recorded at 2 hours (**Figure S18**). Signals in RNP reactions conducted without dNTPs represent RNP-non-specific background responses to lysates in the absence of CATNAP-mediated amplification.

## Discussion

Here, we developed a novel one-pot nucleic acid diagnostic^24^ approach that integrates isothermal linear DNA amplification with CRISPR-Cas12a detection. Our method uses a nicking endonuclease and a strand-displacing polymerase to continuously generate single-stranded DNA targets from a double-stranded template. These ssDNAs activate the *trans*-cleavage activity of a Cas12a-crRNA complex specifically designed to recognize targets adjacent to non-canonical dipurine PAMs, avoiding cleavage of the original template. We demonstrated the system’s efficacy by rapidly detecting HPV-16 DNA, achieving high specificity capable of discriminating between different oncogenic and low-risk HPV strains, and successfully detecting HPV-16 within complex cervical cancer cell lysates, highlighting its potential for point-of-care diagnostics.

The combination of isothermal linear amplification and CRISPR-based detection addresses significant challenges faced by nucleic acid diagnostics in low-resource settings^25,26^. While CRISPR systems offer high specificity, their sensitivity often requires target pre-amplification^1,27^. Conventional exponential amplification methods like PCR, LAMP, and RPA, although sensitive, suffer from drawbacks that limit their point-of-care applicability^28,29^. They often require specialized equipment (thermocyclers for PCR), complex primer design (LAMP), are prone to contamination leading to false positives (especially in settings with limited infrastructure), and can make quantification difficult due to saturation effects^11,30,31^. Our linear amplification approach mitigates contamination risk as it avoids exponential product accumulation, potentially simplifies quantification, and operates isothermally, eliminating the need for thermocycling equipment^11^. The one-pot format further simplifies the workflow, reducing hands-on time and the need for multiple reaction steps, aligning well with the REASSURED criteria for point-of-care diagnostics in resource-limited environments^32,33^.

The limit of detection (LOD) is intrinsically linked to the rate at which activated Cas12a accumulates and cleaves the reporter substrate. This rate depends on both the efficiency of ssDNA production by the nickase/polymerase system and the subsequent efficiency of Cas12a activation by these specific ssDNA molecules generated near nicking sites containing dipurine PAMs. Our strategy aims to maximize this process by enabling continuous ssDNA production without depleting the template DNA, thus fueling sustained *trans*-cleavage activity. The LOD is also limited by background noise, which can arise from non-specific Cas12a activation by off-target sequences (potentially exacerbated in cell lysates) or inherent reporter instability. Further optimization to improve the LOD will involve exploring engineered strand-displacing polymerases with enhanced processivity, refining buffer components for optimal coordinated enzymatic function, and Cas12a protein engineering to boost catalytic rates, enhance activity at room temperature, and minimize non-specific activation^34^.

This study uses a fluorescence-based readout for rapid assay development. However, alternative detection schemes are necessary for low-resource settings. Lateral flow assays (LFAs) offer a simple, equipment-free, and cost-effective visual detection method well-suited for point-of-care applications^35^. CRISPR-Cas systems have already been successfully coupled with LFA strips^36,37^. In these systems, Cas *trans*-cleavage activity releases tagged reporter molecules, which then generate a visible signal on the strip^1,35^. Ongoing work in our laboratories aims to integrate CATNAP-based linear amplification and Cas12a detection with an LFA readout^38^. Future work will also establish the LOD in more challenging clinical matrices such as saliva or urine, which may contain inhibitors and high background DNA/RNA concentrations^39^. Further, streamlining sample preparation and integrating it with the one-pot CATNAP reaction will ultimately result in a rapid, cost-effective, and widespread “sample-to-answer” device^35,40^. Addressing these areas will accelerate the deployment of CRISPR-based diagnostics globally.

## Supporting Information

This article contains Supplemental Figures S1-S18 and Supplemental Tables S1-S10.

## Acknowledgements

We thank members of the Finkelstein laboratory for carefully reading the manuscript.

## Author Contributions

S.S. purified proteins, SS, RK, KR and EN performed biochemical experiments, and analyzed the data. S.S., E.A.N., and I.J.F. prepared figures and wrote the manuscript with input from all co-authors.

## Funding Information

This work was supported by a sponsored research agreement from Global Health Labs, a generous gift from Tito’s Handmade Vodka, a College of Natural Sciences Catalyst award (to I.J.F.), and the Welch Foundation (F-1808 to I.J.F.). The content is solely the responsibility of the authors and does not necessarily represent the official views of the sponsors.

## Conflicts of Interest

The authors declare no competing financial interests.

## Materials and Methods

### Reagents

Nt.BsmAI (cat. no. R0121S), Klenow Fragment (3’→5’ exo-; cat. no. M0212S), dNTPs (cat. no. N0447S), NEBuffer™ 2 (cat. no. B7002S), and rCutSmart™ Buffer (cat. no. B6004S) were purchased from NEB. For all experiments except when comparing orthologs, LbCas12a (cat.no. M0653S) from NEB was used. 1X Cas12 assay buffer (CAB) was composed of 10mM Tris-HCl, pH7.5, 10 mM MgCl_2_, 1.0 mM TCEP, 0.01% IGEPAL CA-630, and 40ug/mlBSA. 858-bp HPV-16 E6E7 gBlock was purchased from IDT. Plasmids encoding HPV types were purchased from ATCC: 16 (cat. No. 45113D); 18 (cat. no. 45152D); 52 (cat. no. VRMC-29™); 6b (cat. no. 45150D).

### Amplification of DNA Targets

Amplification components consisting of 0.13 U/µL Nt.BsmAI, 0.13 U/µL Klenow Fragment(exo-), 50mM dNTPs, and various amounts of template in 1X Cas12 assay buffer (CAB) or rCutSmart, were mixed at room temperature, then incubated at 37°C (unless indicated) for the desired time. Amplification products were quantified by qPCR (**Figure S3**) or used as targets for activating RNP cleavage of *trans*-substrate FQ-C_10_ (**Figure S1**). For “one-pot” reactions, RNP, amplification components, and FQ-C_10_ *trans*-substrate were mixed at room temperature, then incubated at 37°C or room temperature, and reaction products were monitored continuously (**Figure S2**). Control reactions were carried out in amplification reactions lacking dNTPs or templates or in which enzymes were replaced with equivalent volumes of their respective storage buffers.

### Quantification of Amplification Products

100 fM of the 858-bp HPV-16 E6E7 gBlock fragment and dNTPs, diluted into 1X Cas12 assay buffer, were incubated at 37°C with or without nicking enzyme (Nt.BsmAI) and DNA polymerase (Klenow exo-). At the indicated times, aliquots were removed and heat-inactivated by incubation at 80°C for 20 minutes. Quantitative PCR as performed on the aliquots using PrimeTime® Gene Expression Master Mix and PrimeTime Std® qPCR primers and probes (IDT) on a CFX Real-Time PCR Detection System (Bio-Rad) instrument. The cycling protocol consisted of 95°C for 3 min followed by 40 repeats of 95°C for 5 s and 60°C for 60 s (data collection). Linear 858-bp HPV-16 DNA fragment was used as a PCR Standard. The number of copies observed in samples lacking nicking enzyme/polymerase, representing the input material, was subtracted from copies observed in their presence to obtain the number of copies amplified in the reactions.

### One-Pot CATNAP Reactions

One pot CATNAP reaction consists of 0.13 U/µL Nt.BsmAI, 0.13 U/µL Klenow Fragment, 50 mM dNTPs, 2nM Cas12a, 1nM nM crRNA, template (HPV-plasmid), FQ-C_10_ *trans*-substrate and selected 20mM Tris-acetate, 10mM Magnesium acetate, 50mM Potassium acetate, 100µg/ml Albumin pH 7.9.

### Quantification of *trans*-substrate Cleavage Products

Fluorescence measurements were carried out in 384-well plates on an Agilent BioTek Synergy microplate reader (λ_ex_ = 490 nm, 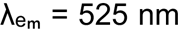). When indicated, fluorescence traces obtained from reactions containing RNP and *trans* substrate but lacking target were subtracted from the corresponding test traces. Resulting fluorescence values were converted to molar concentrations of cleaved *trans*-substrate by interpolation against standard curves generated from dilution series of Cas12a-cleaved substrates.

### Expression and Purification of Cas12a Orthologs

Full length Cas12a orthologs (**Table S10**) were produced in E. coli as N-terminal His_6_,TwinStrep,SUMO fusions and induced with IPTG. Cleared lysates were applied to StrepTactin resin, the SUMO tag removed by protease, and the proteins polished by Superdex 200 size-exclusion chromatography. Peak fractions were pooled, flash-frozen in liquid nitrogen, stored at –80°C, and quantified by Bradford assay using BSA as a standard.

### Preparation of Crude Cell Lysates

Tissue culture cells (C33A, Caski) were grown to confluency in Eagle’s Minimum Essential Medium containing 10% fetal bovine serum, penicillin (100 U/mL), and streptomycin (100 µg/ml). Adherent cells were harvested and resuspended in 10 mM Tris-HCl, pH 7.5, 1.0 mM EDTA. Samples were first heated at 95°C for 20 minutes, then Proteinase K (60 μg/mL) was added. Samples were then heated at 65°C for 20 minutes, then 95°C for 20 minutes, and stored at −80°C until use. DNA content was quantified by Qubit™ 1X dsDNA High Sensitivity Kit (ThermoFisher, cat. no. Q33230). Cell lysates corresponding to 1.0 ng DNA were subjected to “one pot” CATNAP reactions containing 6.3 nM BsCas12a, 3.1 nM crRNA, 100 nM substrate (FQ-C10), 1.25% Nt.BsmAI, 1.25% KF exo-, 50 uM each dNTP, in 1X Cas12 assay buffer at 37°C.

## Notes

### Competing Interest Statement

The authors have declared no competing interest.

