## Supplement for "One-Pot Isothermal Linear Amplification and Cas12a-based Nucleic Acid Detection"

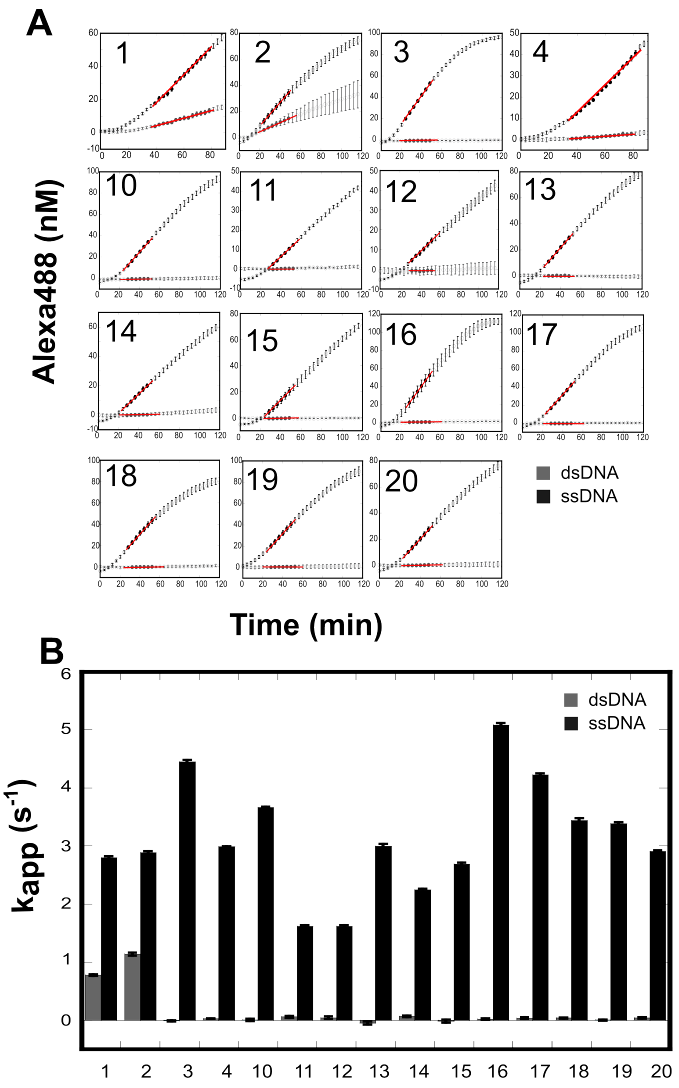


**Figure S1. RNPs using dipurine PAMs are efficiently activated by ssDNA, but not dsDNA targets**.

(A) Time courses for cleavage of *trans*-substrate FQ-$C_{10}$ by RNPs activated by plasmids containing HPV-16 (grey) or 40-nt ssDNA targets (black). Symbols represent the mean background-subtracted product (± SD) of triplicates. Filled symbols represent values in linear ranges used to calculate steady-state cleavage rate taken from slope of solid lines, which, when normalized to target concentration, yields apparent turnover $k_{app}$. (B) Apparent turnover of RNPs activated by dsDNA (grey) and ssDNA (black) targets (bar ± SE) from (A). Summarized in **Table S10**. crRNA sequences are summarized in **Table S1**. Only RNP-1 and 2, which use a dipyrimidine PAM, are efficiently activated by both ssDNA and dsDNA targets.

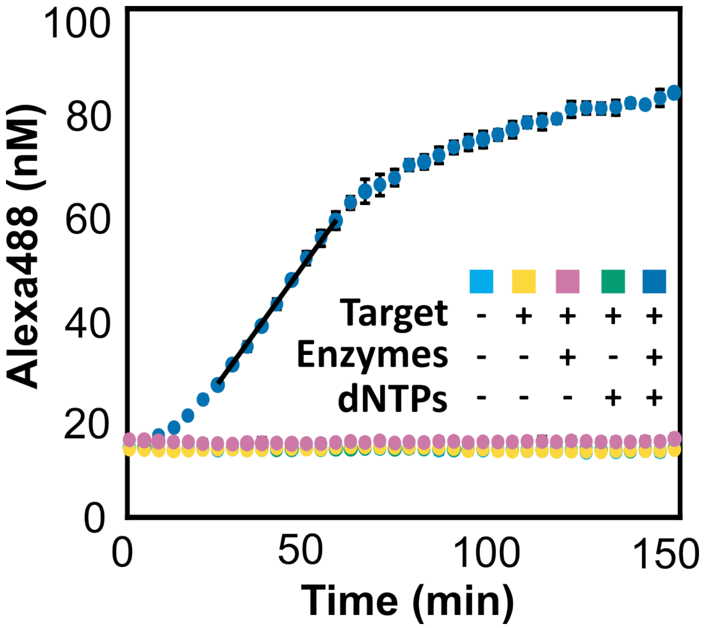


**Figure S2. All enzyme components are required for CATNAP.**

Cleavage of FQ-$C_{10}$ *trans*-substrate by RNP-3 in response to amplification reactions with or without HPV target, nicking enzyme/polymerase, or dNTPs in "one-pot" reactions. Symbols represent mean product (± SD) of triplicates. Values observed at 60 minutes are plotted in **Figure 2C**. Rates of substrate cleavage calculated in linear ranges are provided in **Table S3**.

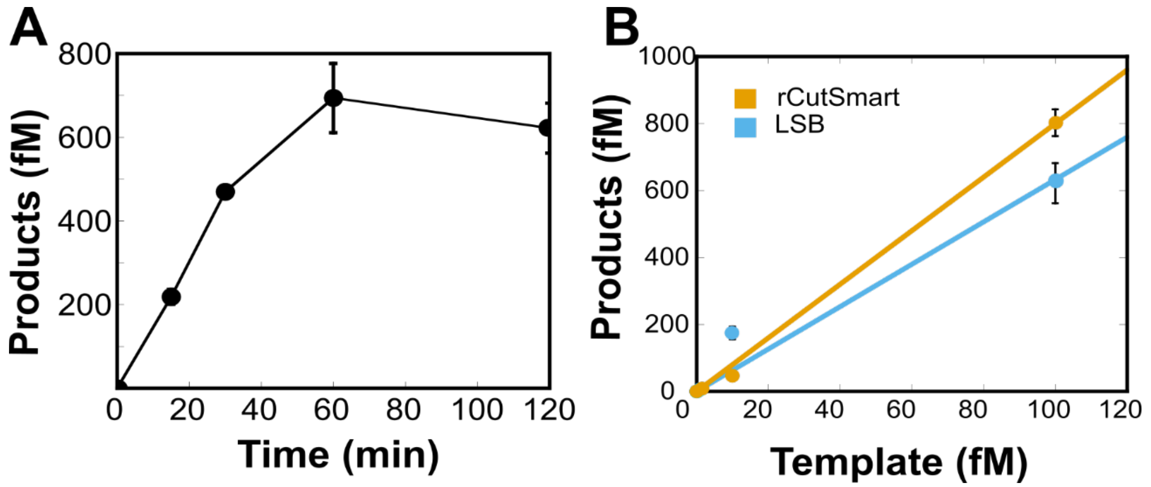


**Figure S3. Quantification of ssDNA synthesized during amplification employing a nicking enzyme and DNA polymerase**.

(A) Amplified reaction products initially accumulate linearly. Symbols represent mean copies (± SD) of triplicates measured for 100 fM input template in amplification reactions containing dNTPs, which was corrected for the number of copies measured from reactions conducted without dNTPs, representing input DNA. After 60 minutes, approximately seven copies of amplified product are generated per input target. (B) Quantification of ssDNA synthesized by amplification enzymes in different reactions buffers. Nicking-polymerase reactions were carried out in Cas12a Assay Buffer (CAB) and rCutSmart Buffer using varied concentrations of dsDNA template. Symbols represent mean copies (± SD) determined by qPCR in triplicate and corrected for input template, measured in control amplification reactions lacking enzymes, and converted to concentration. Linear fitting, indicated by solid line, provides estimation of product synthesis at 5.9 (± 0.8) copies/template (CAB) and 8.1 (± 0.2) copies/template (rCutSmart Buffer).

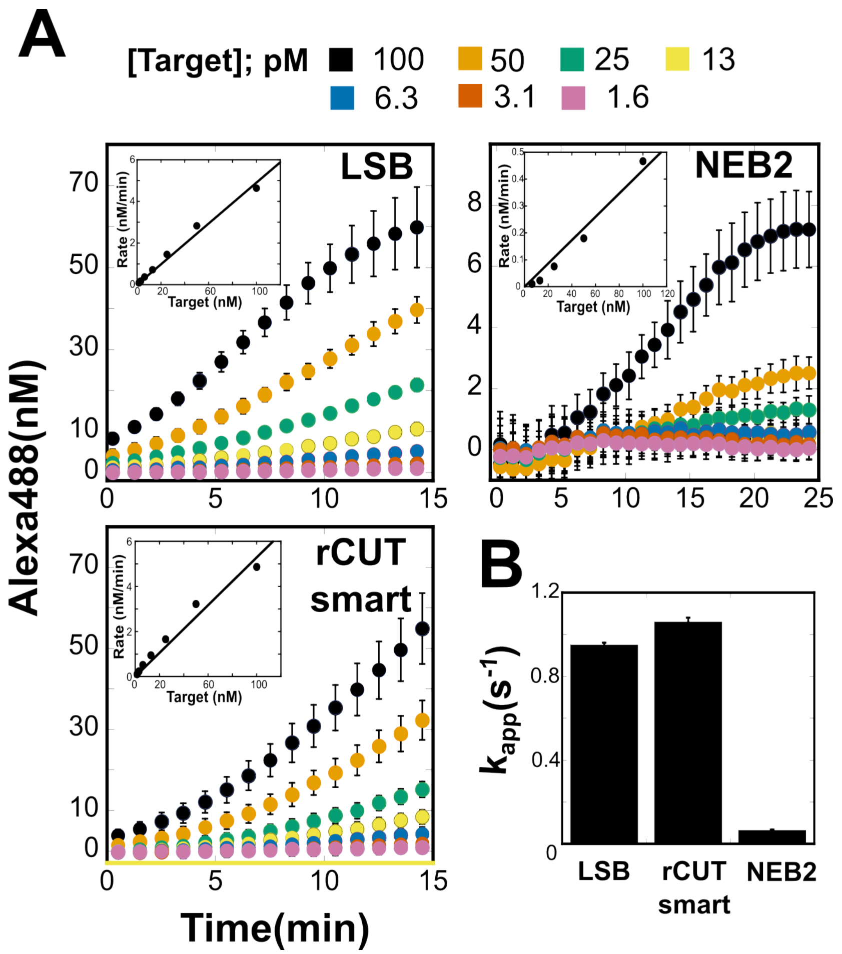


**Figure S4. Buffer optimization for *trans*-substrate cleavage reactions.**

(A) Cleavage of *trans*-substrate FQ-$C_{10}$ by LbCas12a RNP activated with the indicated concentrations of a dsDNA target, performed in reactions containing Cas12a Assay Buffer (CAB), rCutSmart Buffer, and NEBuffer2. Symbols represent mean product (± SD) of triplicates. Cleavage rates were calculated from steady-state portions of the reactions and plotted against target concentration (insets). Linear fitting (solid lines) yielded slopes representing apparent substrate turnover $k_{app}$. (B) Summary of apparent turnover from (A).

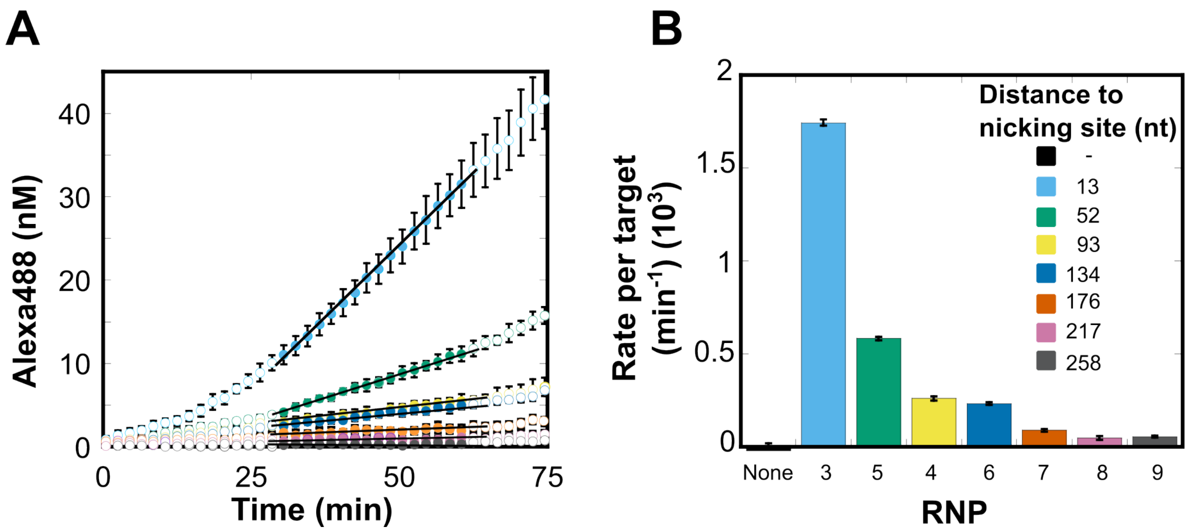


**Figure S5. *Trans*-substrate cleavage by RNPs reacted with amplified reaction products in 2-step reactions**

Cleavage of FQ-$C_{10}$ *trans*-substrate by the indicated CATNAP RNPs activated by products of nicking-polymerase generated in separate reactions. Symbols in time course (A) represent the mean background-corrected product (± SD) of triplicates. Solid lines and symbols represent linear ranges, from which cleavage rates (± SE) were calculated in (B). The background rate measured for Cas12a in the absence of crRNA (None) was -0.001 (±0.001) nM $min^{-1}$.

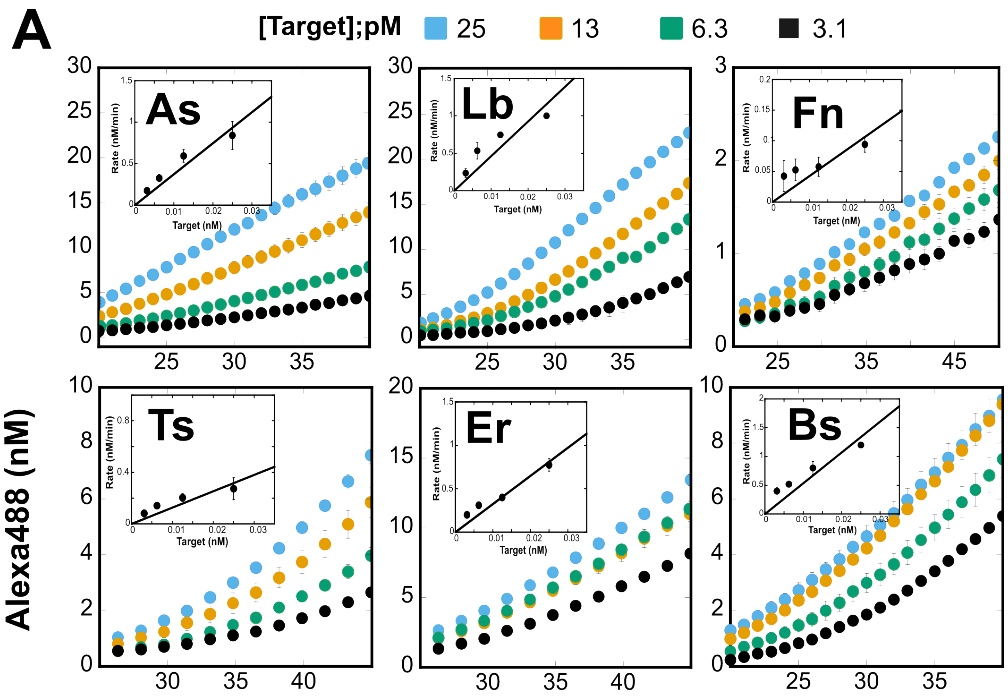


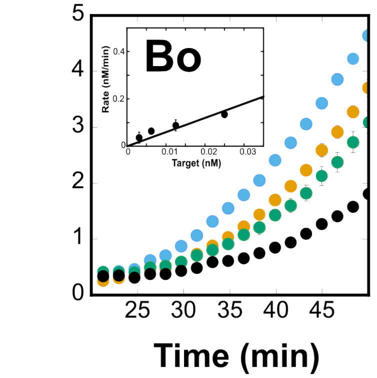


**B**

| Cas12a ortholog | Rate per target ($min^{-1}$) |
|---|---|
| Lb | 38 (± 5) |
| As | 46 (± 4) |
| Fn | 3 (± 1) |
| Er | 23 (± 4) |
| Ts | 13 (± 3) |
| Bs | 54 (± 1) |
| Bo | 6 (± 1) |

**Figure S6. Impact of Cas12a orthologs on CATNAP *trans*-cleavage rates.**

(A) Time course of CATNAP *trans*-cleavage of FQ-$C_{10}$, where symbols represent mean product (± SD) of triplicates. Reactions were performed using 2.0 nM RNP (3) in rCutSmart buffer with varying concentrations of HPV-16 plasmid as template. Cas12a orthologs include: **As,** *Acidaminococcus sp.* Cas12a (AsCas12a); **Lb,** *Lachnospiraceae bacterium* Cas12a (LbCas12a); **Fn**, *Francisella novicida* Cas12a (FnCas12a); **Er,** *Eubacterium rectale* Cas12a (ErCas12a); **Ts,** *Thiomicrospira sp.* Cas12a (TsCas12a); **Bs,** *Butyrivibrio sp.* Cas12a (BsCas12a); **Bo**, *Bacteroidetes oral* taxon Cas12a (BoCas12a). Inset: rate of substrate cleavage (mean ± SE) calculated during linear phases of time course, between 25–35 or 30–40 minutes, plotted against input target concentration. Solid lines represent linear fitting, yielding slopes corresponding to cleavage rate per input target. (B) CATNAP *trans*-substrate cleavage rates normalized to input target concentration, taken from insets in (A), plotted in **Figure 3A**. BsCas12a represents the optimal Cas12a CATNAP RNP under these conditions.

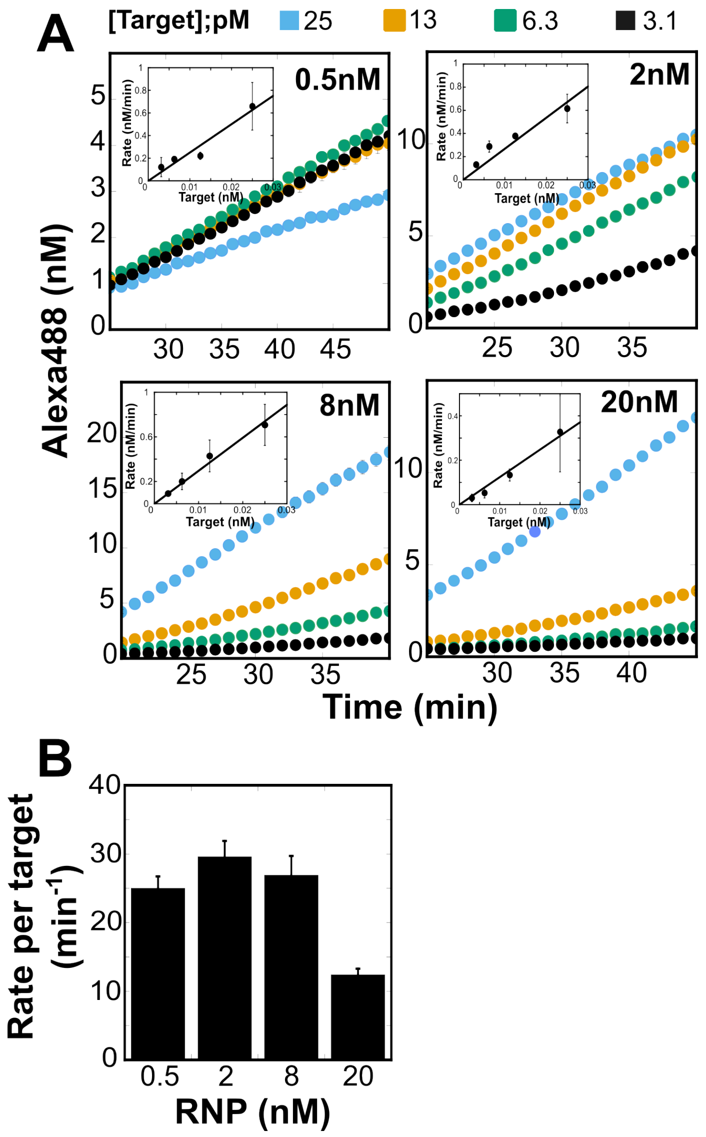


**Figure S7. Optimizing RNP concentrations for CATNAP.**

(A) Time course of CATNAP *trans*-cleavage of FQ-$C_{10}$, where symbols represent mean product (± SD) of triplicates. Reactions were performed using varying concentrations of LbCas12a RNP (3) in rCutSmart buffer with varying concentrations of HPV-16 plasmid as template. Inset: rate of substrate cleavage (mean ± SE) calculated during linear phases of time course, between 25–35 or 30–40 minutes, plotted against input target concentration. Solid lines represent linear fitting, yielding slopes corresponding to cleavage rate per input target. (B, C) CATNAP *trans*-substrate cleavage rates (± SE) normalized to input target concentration, taken from insets of (A). 2.0 nM represents the optimal RNP concentration for CATNAP.

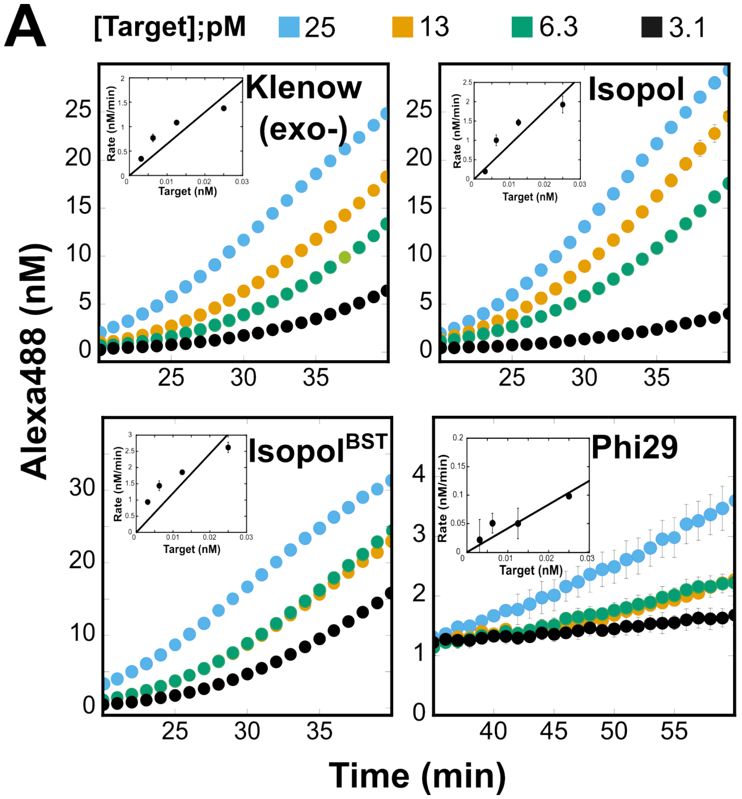


**B**

| Cas12a ortholog | Rate per target ($min^{-1}$) |
|---|---|
| **Klenow Exo (-)** | 88.0 (± 9.8) |
| **IsoPol** | 64.2 (± 6.9) |
| **IsoPol $^{BST+}$** | 121.0 (± 13.4) |
| **Phi29** | 4.2 (± 0.6) |

**Figure S8. Evaluating different DNA polymerases for CATNAP.**

(A) Time course of CATNAP *trans*-cleavage of FQ-$C_{10}$, where symbols represent mean product (± SD) of triplicates. Reactions were performed using 2.0 nM of LbCas12a RNP (3) in rCutSmart buffer with different DNAP and varying concentrations of HPV-16 plasmid as template. Inset: rate of substrate cleavage (mean ± SE) calculated during linear phases of time course, between 25–35 or 40–50 minutes, plotted against input target concentration. Solid lines represent linear fitting, yielding slopes corresponding to cleavage rate per input target. (B) CATNAP *trans*-substrate cleavage rates (± SE) normalized to input target concentration, taken from insets of panel A. Values are plotted in **Figure 3B**. IsoPol$^{BST+}$ generated the largest CATNAP signal.

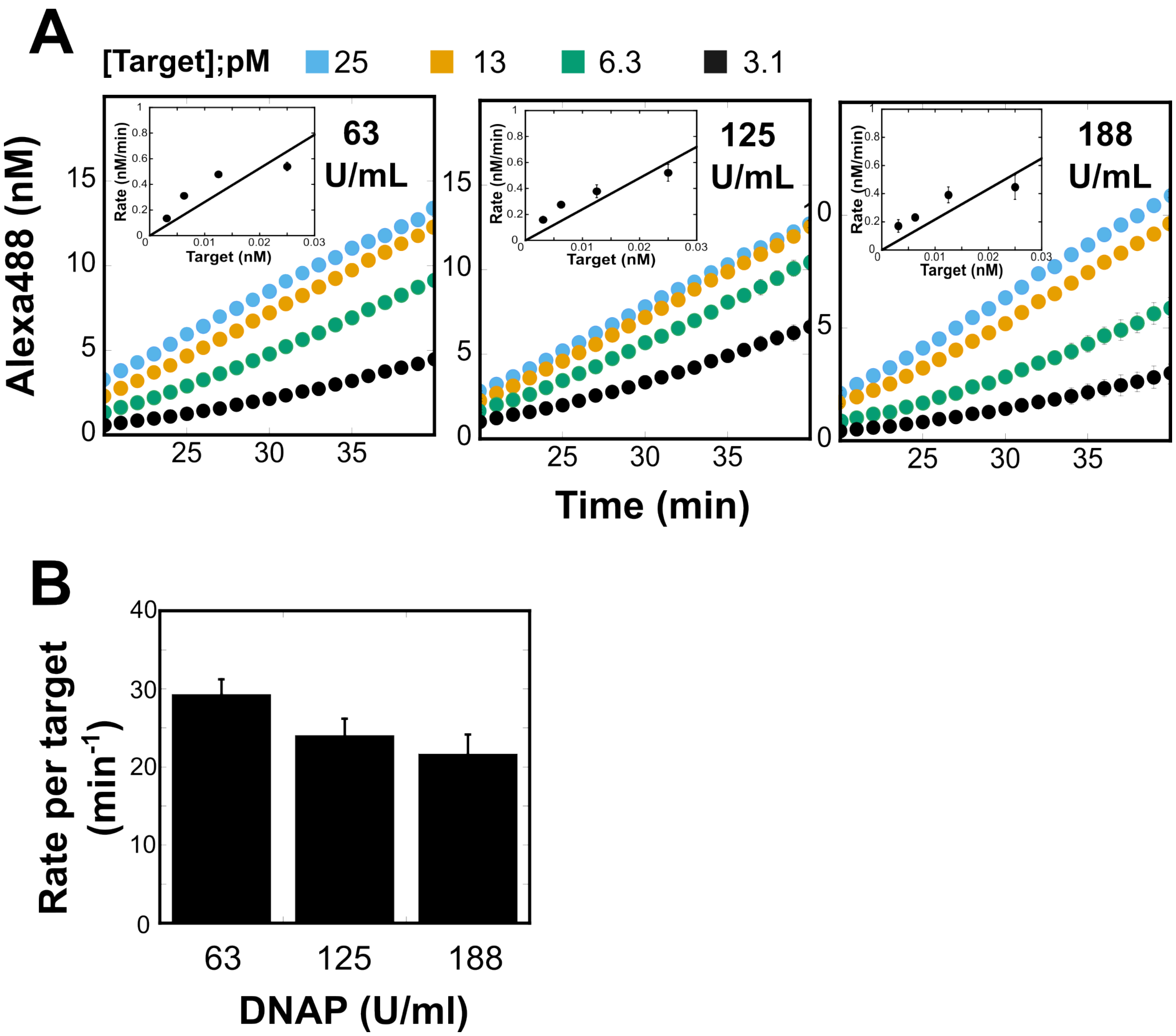


**Figure S9. Optimization of Klenow (exo-) DNA polymerase concentration in CATNAP reactions.**

(A) Time course of CATNAP *trans*-cleavage of FQ-$C_{10}$, where symbols represent mean product (± SD) of triplicates. Reactions were performed using 2.0 nM of LbCas12a RNP (3) in rCutSmart buffer with varying concentrations of Klenow (exo-) and varying concentrations of HPV-16 plasmid as template. U = activity units as defined by the manufacturer. Inset: rate of substrate cleavage (mean ± SE) calculated during linear phases of time course, between 25–35 or 30–40 minutes, plotted against input target concentration. Solid lines represent linear fitting, yielding slopes corresponding to cleavage rate per input target. (B) CATNAP *trans*-substrate cleavage rates (± SE) normalized to input target concentration, taken from insets of (A).

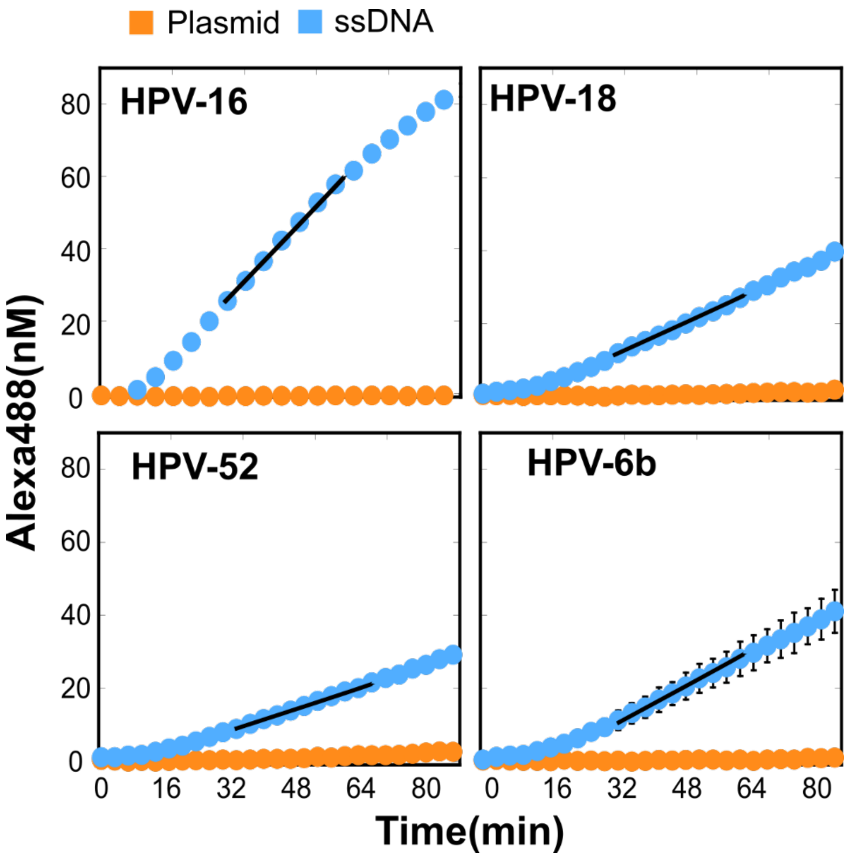


**Figure S10. Development of RNPs for HPV type-specific CATNAP**.

Time course for cleavage of *trans*-substrate FQ-$C_{10}$ by RNPs specific to HPV-16, -18, -52, and -6b, activated by short, 39–40-nt ssDNA targets (blue) or plasmid DNA encoding the corresponding HPV genomes (orange). Symbols represent mean background-subtracted product (± SD) of triplicates. Linear fitting (solid lines) yielded slopes corresponding to steady-state cleavage rates, which, when normalized to target concentration, yield apparent turnover $k_{app}$, provided in Table S8. dsDNA plasmids on their own do not activate Cas12a CATNAP RNPs.

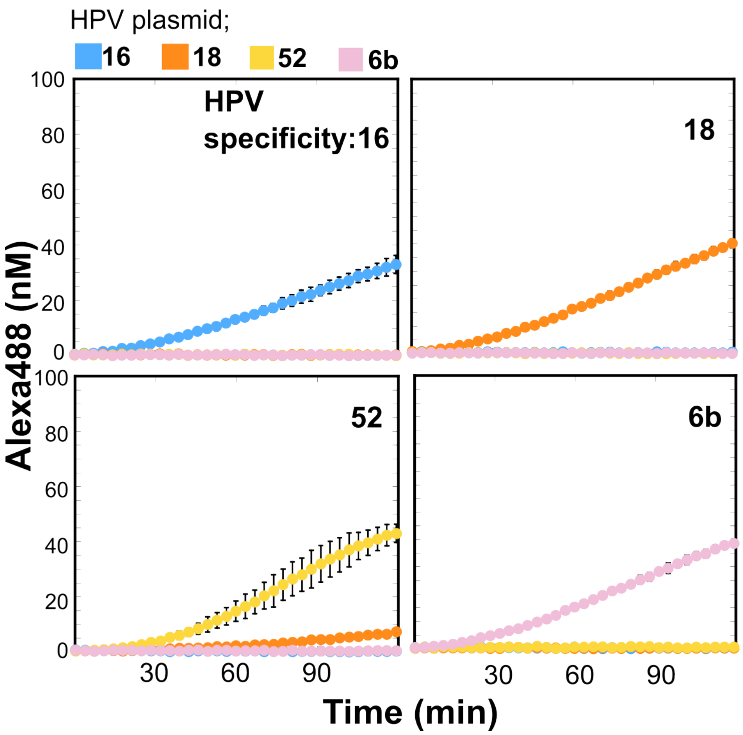


**Figure S11. Type-specific HPV detection by CATNAP.**

CATNAP reactions were performed at 25°C with RNPs designed to HPV-16, -18, -52, and -6b (**Figure S10**) using plasmids containing each type. Symbols represent mean (± SE) of background-corrected product measured in triplicate.

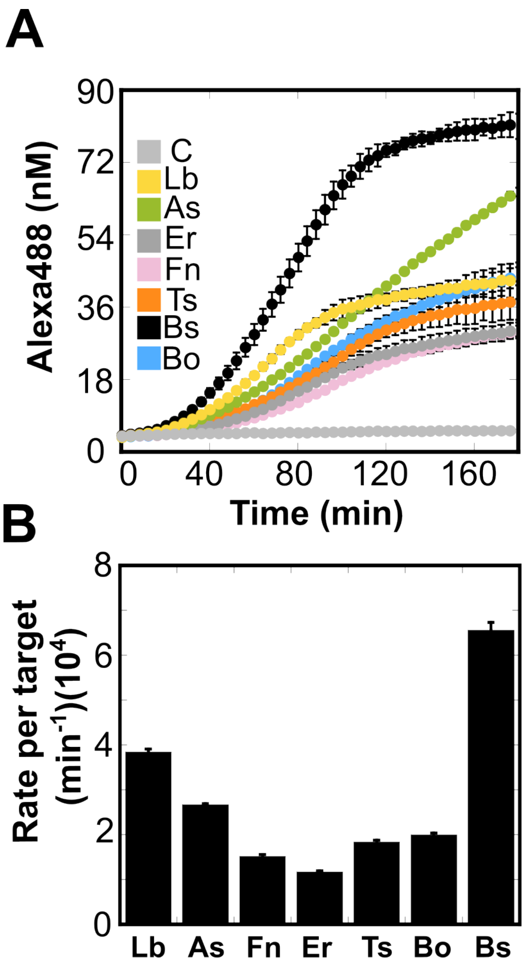


**Figure S12. Cas12a ortholog optimization for CATNAP at 25°C.**

(A) Time course for CATNAP cleavage of *trans*-substrate FQ-$C_{10}$ measured using 2.0 nM RNP, 100 nM FQ-$C_{10}$ *trans*-substrate, and rCutSmart buffer at 25°C. Control (C) shows no RNP. Symbols represent mean background-corrected product (± SD) of triplicates. (B) Apparent *trans*-cleavage rates (mean ± SE, n = 3) obtained from slopes of linear portions of time courses in (A). Ortholog abbreviations: C, control; Lb, *Lachnospiraceae bacterium* Cas12a; As, *Acidaminococcus sp.* Cas12a; Fn, *Francisella novicida* Cas12a; Er, *Eubacterium rectale* Cas12a; Ts, *Thiomicrospira sp.* Cas12a; Bs, *Butyrivibrio sp.* Cas12a; Bo, *Bacteroidetes oral* taxon Cas12a.

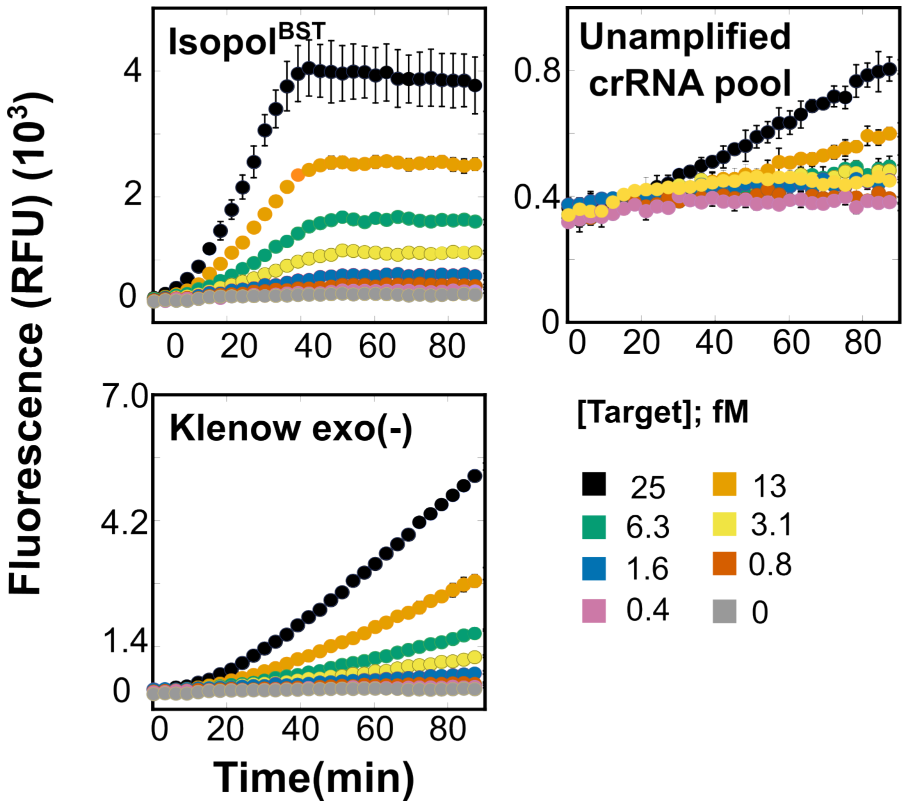


**Figure S13. Comparison of HPV-16 plasmid detection between CATNAP reactions using two different DNA polymerases and amplification-free reactions using RNP pools.**

Time course of *trans*-cleavage of FQ-$C_{10}$ generated in response to varying concentrations of HPV-16 plasmid, where symbols represent mean fluorescence of product (± SD) of triplicates. CATNAP (left panels) contained 2.0 nM BsCas12a RNP (pools of 12 CATNAP RNPs) and either IsoPol$^{BST+}$ or Klenow (exo-) in rCutSmart buffer. Unamplified reactions (right panel) contained 2.0 nM BsCas12a RNP (pools of 20 non-CATNAP RNPs) in CAB. LOD and FOM were calculated at each time point (**Figure 5**).

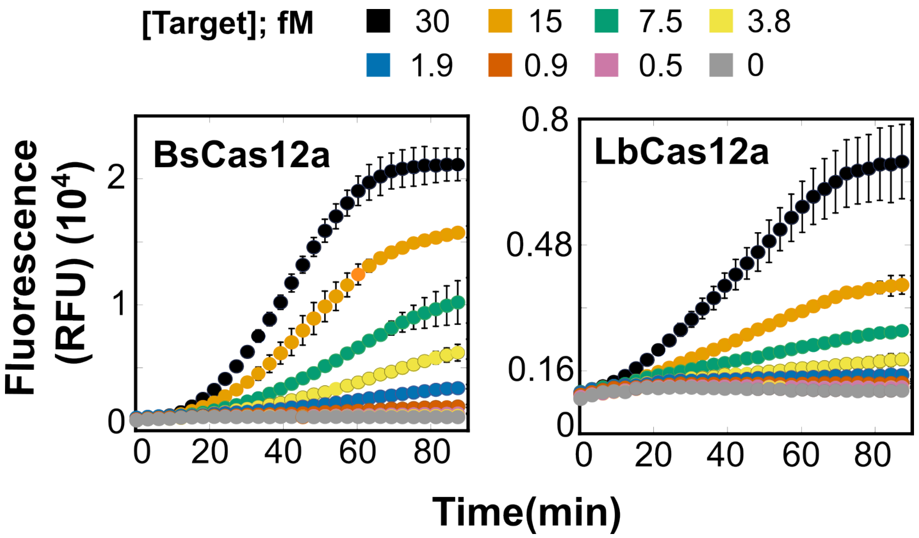


**Figure S14. Comparison of CATNAP reactions for detection of HPV-16 plasmid using two different Cas12a orthologs.**

Time course of CATNAP *trans*-cleavage of FQ-$C_{10}$, where symbols represent mean fluorescence of product (± SD) of triplicates. Reactions were performed using 2.0 nM of Bs or LbCas12a RNP (pools of 12 RNPs) in rCutSmart buffer with varying concentrations of HPV-16 plasmid as template. LODs and FOMs were calculated at each time point (**Figure 5**).

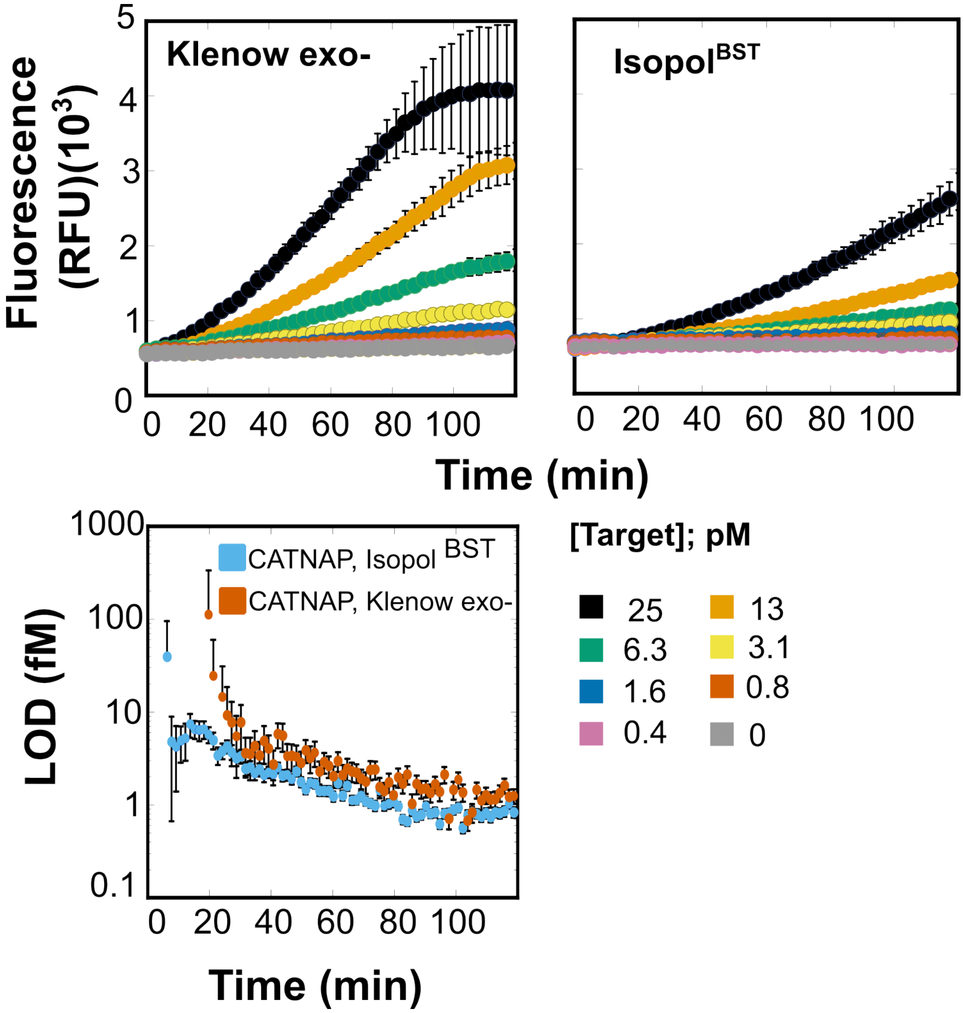


**Figure S15. CATNAP reporter cleavage kinetics and LOD determination at 25°C.**

(A) Time course of CATNAP *trans*-cleavage of FQ-$C_{10}$ at 25°C, where symbols represent mean fluorescence of product (± SD) of triplicates. CATNAP reactions were performed using 2.0 nM of BsCas12a RNP (pools of 12 RNPs) in rCutSmart buffer with varying concentrations of HPV-16 plasmid as template. (B) Assay sensitivity, where symbols represent mean LOD (± SE) calculated at each time point in (A)

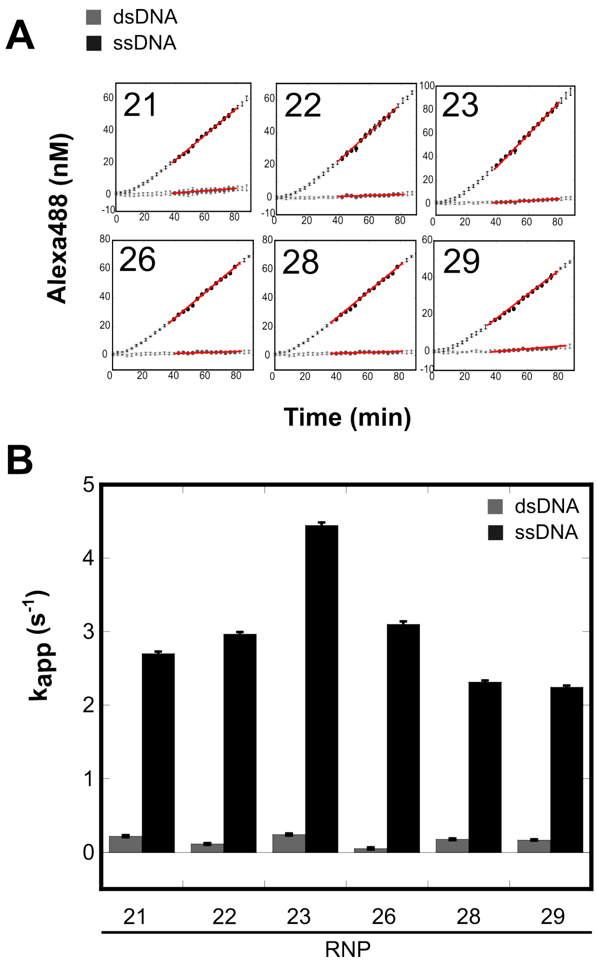


**Figure S16. RNPs using dipurine PAMs are efficiently activated by ssDNA but not by dsDNA targets.** A) Time courses for cleavage of *trans*-substrate FQ-$C_{10}$ by RNPs activated by plasmids containing HPV-16 (grey) or 40-nt ssDNA targets (black). Symbols represent mean background-subtracted product (± SD) of triplicates. Filled symbols represent values in linear ranges used to calculate steady-state cleavage rate taken from slope of solid lines, which, when normalized to target concentration, yields apparent turnover $k_{app}$. (B) Apparent turnover of RNPs activated by dsDNA (grey) and ssDNA (black) targets (bar ± SE) from (A). crRNA sequences are summarized in **Table S1.**

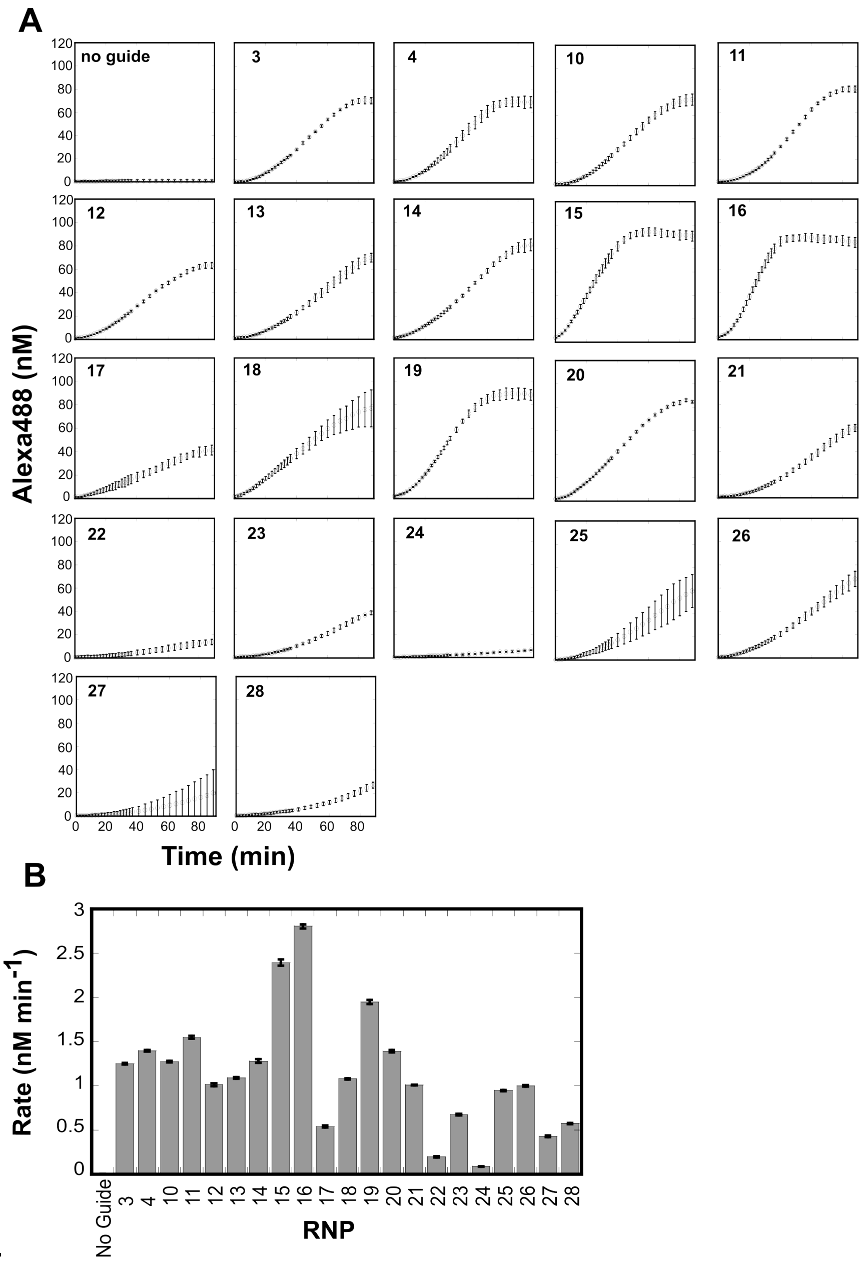


**Figure S17. Screening HPV-16 CATNAP guides.**

(A) CATNAP reactions were performed with HPV-16-specific RNPs using FQ-$C_{10}$ and HPV-16 plasmid as input target. Symbols represent the mean background-subtracted product (± SD) of triplicates. Slopes of linear portions were calculated to obtain *trans*-cleavage rate. (B) Results of guide screening. Bars (± SE) represent *trans*-cleavage rates from (A) normalized to input DNA. crRNA sequences are summarized in **Table S1**. RNP-16 generates the greatest CATNAP signal.

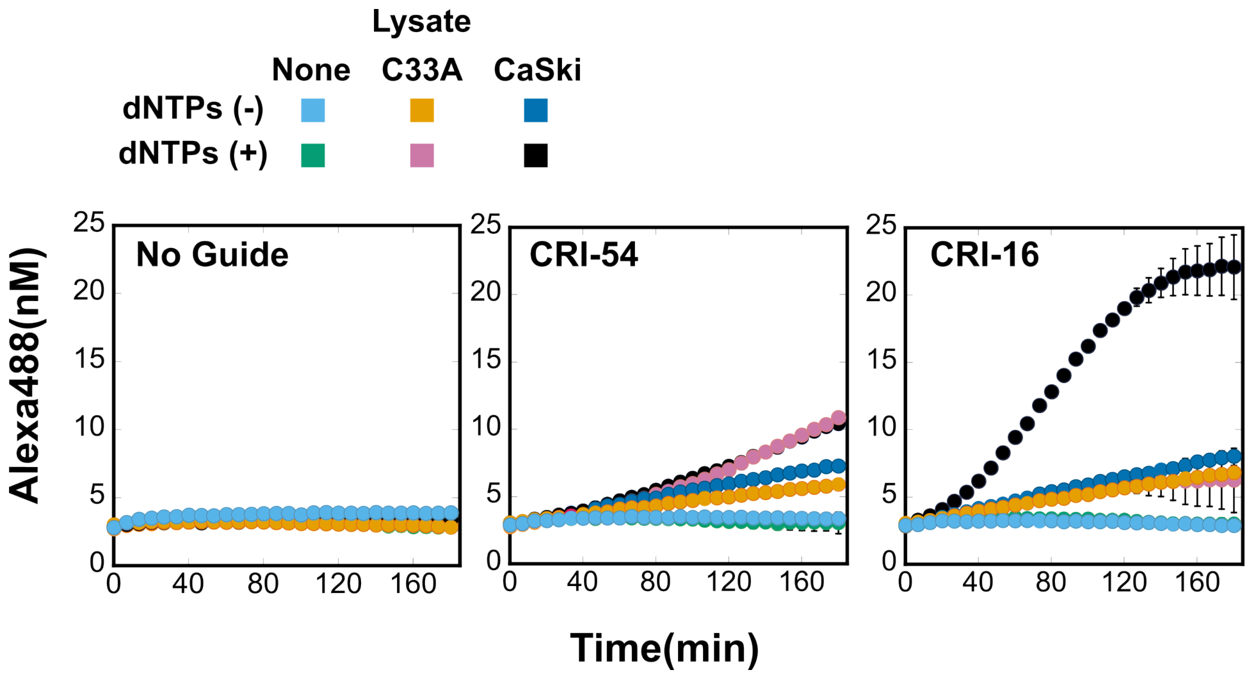


**Figure S18. CATNAP detection of HPV-16 in crude cell lysates.**

Time courses of CATNAP reactions conducted on crude cell lysates using HPV-16-specific CATNAP RNP (CRI-16) and control conditions (no RNP and non-CATNAP RNP-54 specific to *M. ulcerans*). Symbols represent mean product (± SD) of triplicates. Values recorded at 2 hours are plotted in **Figure 5**. Reactions conducted without dNTPs represent background responses to lysates in the absence of CATNAP-mediated amplification.

### Supplemental Tables

| Table S1. Sequences of short synthetic nucleic acids | |
|---|---|
| **crRNA specific to HPV-16 (non-CATNAP)** | |
| RNP | Sequence |
| 2 | UAAUUUCUACUAAGUGUAGAU**CAUAAUAUAAGGGGUCGGUG** |
| 29 | UAAUUUCUACUAAGUGUAGAU**AACCGAAACCGGUUAGUAUA** |
| 30 | UAAUUUCUACUAAGUGUAGAU**AUGUAUAAAACUAAGGGCGU** |
| 31 | UAAUUUCUACUAAGUGUAGAU**UGCACAGAGCUGCAAACAAC** |
| 32 | UAAUUUCUACUAAGUGUAGAU**UAUGCACCAAAAGAGAACUG** |
| 33 | UAAUUUCUACUAAGUGUAGAU**AGGACCCACAGGAGCGACCC** |
| 34 | UAAUUUCUACUAAGUGUAGAU**AUGAUCUGCAACAAGACAUA** |
| 35 | UAAUUUCUACUAAGUGUAGAU**CCUCUGAGCUGUCAUUUAAU** |
| 36 | UAAUUUCUACUAAGUGUAGAU**CUGCGACGUGAGGUAUAUGA** |
| 37 | UAAUUUCUACUAAGUGUAGAU**AUAUGCUGUAUGUGAUAAAU** |
| 38 | UAAUUUCUACUAAGUGUAGAU**GGGAUUUAUGCAUAGUAUAU** |
| 39 | UAAUUUCUACUAAGUGUAGAU**GGUUGUGCGUACAAAGCACA** |
| 40 | UAAUUUCUACUAAGUGUAGAU**AUGGGCACACUAGGAAUUGU** |
| 41 | UAAUUUCUACUAAGUGUAGAU**UAAUAUUAUAUCAUGUAUAG** |
| 42 | UAAUUUCUACUAAGUGUAGAU**CAUACAAACUAUAACAAUAA** |
| 43 | UAAUUUCUACUAAGUGUAGAU**UGCAUGAUUACAGCUGGGUU** |
| 44 | UAAUUUCUACUAAGUGUAGAU**CAACCAGAGACAACUGAUCU** |
| 45 | UAAUUUCUACUAAGUGUAGAU**GAACAGCAAUACAACAAACC** |
| 46 | UAAUUUCUACUAAGUGUAGAU**UAACCUUUUGUUGCAAGUGU** |

| | |
|---|---|
| 47 | UAAUUUCUACUAAGUGUAGAU**GAAGAAAAGCAAAGACAUCU** |
| 48 | UAAUUUCUACUAAGUGUAGAU**GUUAAUUAGGUGUAUUAACU** |
| 49 | UAAUUUCUACUAAGUGUAGAU**AGCUGGACAAGCAGAACCGG** |
| 50 | UAAUUUCUACUAAGUGUAGAU**ACAUUGCAUGAAUAUAUGUU** |
| | * Present in 20-RNP pool (29, 30, 31, 32, 33, 34, 35, 36, 37, 38, 39, 40, 42, 43, 44, 45, 46, 47, 48, 50). |
| **crRNA specific to HPV-16 (CATNAP)** | |
| RNP | Sequence |
| 1 | UAAUUUCUACUAAGUGUAGAU**GGGAGUGGUUACAAAAGCAG** |
| 3 | UAAUUUCUACUAAGUGUAGAU**ACACCUACAUUGCAUGAAUA** |
| 4 | UAAUUUCUACUAAGUGUAGAU**CGGUGGACCGGUCGAUGUAU** |
| 5 | UAAUUUCUACUAAGUGUAGAU**AACACGUAGAGAAACCCAGC** |
| 6 | UAAUUUCUACUAAGUGUAGAU**GACAUCUGGACAAAAAGCAA** |
| 7 | UAAUUUCUACUAAGUGUAGAU**GUAUUAACUGUCAAAAGCCA** |
| 8 | UAAUUUCUACUAAGUGUAGAU**ACAGCAAUACAACAAACCGU** |
| 9 | UAAUUUCUACUAAGUGUAGAU**UAUAGACAUUAUUGUUAUAG** |
| 10 | UAAUUUCUACUAAGUGUAGAU**AAUGAUUAUUUAACACAGGC** |
| 11 | UAAUUUCUACUAAGUGUAGAU**AGAUGUUACAGGUAGAAGGG** |
| 12 | UAAUUUCUACUAAGUGUAGAU**AGGUAUAUCAAAUAUUAGUG** |
| 13 | UAAUUUCUACUAAGUGUAGAU**UUGCACGAGGACGAGGACAA** |
| 14 | UAAUUUCUACUAAGUGUAGAU**UGGAUAUACAGUGGAAGUGC** |
| 15 | UAAUUUCUACUAAGUGUAGAU**CCCUGCCACACCACUAAGUU** |
| 16 | UAAUUUCUACUAAGUGUAGAU**UUUGGAGGCUCUAUCAUCAU** |
| 17 | UAAUUUCUACUAAGUGUAGAU**ACUGUAAUAGUUUUUGGUAU** |
| 18 | UAAUUUCUACUAAGUGUAGAU**UAUAUAAUCCUAGGCGUGCC** |
| 19 | UAAUUUCUACUAAGUGUAGAU**CACCUGCAUCAGCAAUAAUU** |

| | |
|---|---|
| 20 | UAAUUUCUACUAAGUGUAGAU**AGACAGUGGCCUCACUAGGC** |
| 21 | UAAUUUCUACUAAGUGUAGAU**GAGAACGAAAAUGACAGUGA** |
| 22 | UAAUUUCUACUAAGUGUAGAU**GAGAUUAUUUGAAAGCGAAG** |
| 23 | UAAUUUCUACUAAGUGUAGAU**CUUAAUGAUAAGAACUGGAA** |
| 24 | UAAUUUCUACUAAGUGUAGAU**ACGUUAGCCUUGAAGUGUAU** |
| 25 | UAAUUUCUACUAAGUGUAGAU**ACCGAAGAAACACAGACGAC** |
| 26 | UAAUUUCUACUAAGUGUAGAU**AAUGUUGUGGAUGCAGUAUC** |
| 27 | UAAUUUCUACUAAGUGUAGAU**GUUUACGUCGUUUUCGUAAC** |
| 28 | UAAUUUCUACUAAGUGUAGAU**AUAUUCAUCCGUGCUUACAA** |
| | * Present in 12-RNP CATNAP pool (3,10-20) |
| **crRNA specific to other HPV types (CATNAP)** | |
| RNP | Sequence |
| 50 | UAAUUUCUACUAAGUGUAGAU**UAAGACAUUAUUCAGACUCU** |
| 51 | UAAUUUCUACUAAGUGUAGAU**UGCCUGCAUAAUAAUAGAUG** |
| 52 | UAAUUUCUACUAAGUGUAGAU**CUGUGCAUGCCCAUACUAAA** |
| **crRNA specific to other targets (non-CATNAP)** | |
| RNP | Sequence |
| **53** | UAAUUUCUACUAAGUGUAGAU**CAUGCCAAUGCGCGACAUUC** |
| 54 | UAAUUUCUACUAAGUGUAGAU**CGCGCCAAAGCGACAGCCAC** |
| **ssDNA targets containing protospacers from HPV-16 recognized by RNP (non-CATNAP)** | |
| Number | Sequence |
| 55 | CTGCTTT**TATACTAACCGGTTTCGGTT**CAACCGATTTCGG |
| 56 | ACCGGTC**CACCGACCCCTTATATTATG**GAATCTTTGCTTT |
| **ssDNA targets containing protospacers from HPV-16 recognized by RNP (CATNAP)** | |

| Number | Sequence |
|---|---|
| 57 | ATACAAA**CTATAACAATAATGTCTATA**CTCACTAATTTTA |
| 58 | GGAATCT**TTGCTTTTTGTCCAGATGTC**TTTGCTTTTCTTC |
| 59 | TCACACA**ACGGTTTGTTGTATTGCTGT**TCTAATGTTGTTC |
| 60 | TAACATA**TATTCATGCAATGTAGGTGT**ATCTCCATGCATG |
| 61 | GACACAG**TGGCTTTTGACAGTTAATAC**ACCTAATTAACAA |
| 62 | ACAAGAC**ATACATCGACCGGTCCACCG**ACCCCTTATATTA |
| 63 | GATTACA**GCTGGGTTTCTCTACGTGTT**CTTGATGATCTGC |
| 64 | GTTTCT**GCCTGTGTTAAATAATCATT**ATCATTTACTATA |
| 65 | TGGCG**CCCTTCTACCTGTAACATCT**GCTGAGTTTCCAC |
| 66 | TACACTT**CACTAATATTTGATATACCT**GTTTTATACCAAT |
| 67 | GTTTTCC**TTGTCCTCGTCCTCGTGCAA**ACTTAATCTGGAC |
| 68 | TCAAACT**GCACTTCCACTGTATATCCA**TGTTTTTTTATAC |
| 69 | GTGCAAC**AACTTAGTGGTGTGGCAGGG**GTTTCCGGTGTCT |
| 70 | AATGTGT**ATGATGATAGAGCCTCCAAA**ATTGCGTAGTACA |
| 71 | AGTTAAA**ATACCAAAAACTATTACAGT**GTCTACTGGATTT |
| 72 | CCCAGT**GGCACGCCTAGGATTATATA**GTCGCACAACACA |
| 73 | AATATAC**AATTATTGCTGATGCAGGTG**ACTTTTATTTACA |
| 74 | TTGGCTGCCTAGTGAGGCCACTGTCTACTTGCCTCCTGT |
| 75 | TCTAAGGTTGTAAGCACGGATGAATATGTTGCACGCACAA |
| 76 | ATTACATGTTACGAAAACGACGTAAACGTTTACCATATTT |
| 77 | GTAGACCCTGCTTTTGTAACCACTCCCACTAAACTTATTA |
| 78 | AAATCTTGATACTGCATCCACAACATTACTGGCGTGCTTT |
| 79 | CTGGATAGTCGTCTGTGTTTCTTCGGTGCCCAAGGCGACG |
| 80 | CAGTTAAATACACTTCAAGGCTAACGTCTTGTAATGTCCA |
| 81 | AAAGGATTTCCAGTTCTTATCATTAAGCTCATACACTGGA |

| | |
|---|---|
| 82 | CCGCTGTCTTCGCTTTCAAATAATCTCCTTTTTGCAGCTC |
| 83 | ACCTGTATCACTGTCATTTTCGTTCTCGTCATCTGATATA |
| **ssDNA targets containing protospacers from other HPV types recognized by RNP (CATNAP)** | |
| Number | Sequence |
| HPV-18 (84) | ATACACA**GAGTCTGAATAATGTCTTAA**TTCTCTAATTCT |
| HPV-52 (85) | GCACAAG**CATCTATTATTATGCAGGCA**GTTCTCGATTACT |
| HPV-6b (86) | CACAACG**TTTAGTATGGGCATGCACAG**GCCTAGAGGTGGG |
| HPV-52 (87) | UAAUUUCUACUAAGUGUAGAU**CGUUGGACAGGGCGCUGUUC** |
| **ssDNA targets containing protospacers from other targets recognized by RNP (non- CATNAP)** | |
| Number | Sequence |
| 88 | TTCTTCG**GAATGTCGCGCATTGGCATG**GAAGTCACACCTT |
| **ssDNA probes and primers for qPCR** | |
| Type | Sequence |
| Forward primer | CCGGTCGATGTATGTCTTGTT |
| Reverse primer | TCATGCAATGTAGGTGTATCTCC |
| Probe | FAM-CAAGAACAC-Zen-GTAGAGAAACCCAGCTGT-IABkFQ |
| Sequences are shown in 5' to 3' direction. For dsDNA, only the TS is shown. Spacer and protospacer sequences are highlighted in bold. Abbreviations: A488, Alexa488; IaBkQ, Iowa Black$^{TM}$ quencher; Zen, ZEN$^{TM}$ quencher; FAM, fluorescein derivative. | |

**Table S2. Cleavage of *trans*-substrate FQ-$C_{10}$ by RNPs activated by dsDNA and ssDNA.**

Apparent turnover $k_{app}$ (± SE) of *trans*-substrate FQ-$C_{10}$ during linear portions of time courses (**Figure 2B**) for HPV-16-specific RNPs reacted with protospacers within long, 858-bp dsDNA (HPV-16 plasmid part) or short, 40-nt ssDNA. Values are plotted in **Figure S1.** RNPs utilizing dipurine PAMs are efficiently activated by ssDNA but not by dsDNA targets.

| RNP | PAM | $k_{app}$ ($s^{-1}$) | |
|---|---|---|---|
| | | Long dsDNA target | Short ssDNA target |
| 1 | GTTG | 0.78(± 0.01) | 2.80 (± 0.03) |
| 2 | ATTC | 1.14 (± 0.03) | 2.88 (± 0.02) |
| 3 | AGAT | -0.01 (± 0.01) | 4.44 (± 0.04) |
| 4 | GGGT | 0.03 (± 0.01) | 2.98 (± 0.01) |

**Table S3. CATNAP using RNPs targeting dipurine and dipyrimidine PAMs.**

CATNAP reactions were performed using different RNPs on the HPV-16 plasmid template, in the presence or absence of dNTPs. Values represent *trans*-substrate FQ-$C_{10}$ cleavage rates (± SE) from linear portions of time courses in **Figure S2**. Values were normalized to input target concentration and plotted in **Figure 2D.** Central dipurines and dipyrimidines of PAMs are underlined.

| RNP | PAM | -dNTP Cleavage rate (nM $min^{-1}$) | +dNTP Cleavage rate (nM $min^{-1}$) |
|---|---|---|---|
| None | - | 0.002 (± 0.001) | 0.007 (± 0.001) |
| 1 | G<u>TT</u>G | 0.055 (± 0.001) | 0.076 (± 0.002) |
| 2 | A<u>TT</u>C | 0.025 (± 0.001) | 0.081 (± 0.001) |
| 3 | A<u>GA</u>T | 0.006 (± 0.001) | 0.316 (± 0.002) |
| 4 | G<u>GG</u>T | 0.007 (± 0.001) | 0.267 (± 0.001) |

**Table S4. Component requirements for CATNAP.**

Rates of *trans*-substrate cleavage (values ± SE) during linear portions of time courses of CATNAP reactions (**Figure S2**). Conditions: 37°C; 1.0 nM RNP-62, 100 nM FQ-$C_{10}$ *trans*-substrate, in the presence or absence of the indicated amplification components and 0.5 pM HPV-16 plasmid.

| | **Amplification component** | | **Cleavage rate (nM $min^{-1}$)** |
|---|---|---|---|
| **Template** | **Enzymes** | **dNTPs** | |
| – | – | – | 0.004 (± 0.005) |
| + | – | – | 0.010 (± 0.005) |
| + | + | – | 0.002 (± 0.003) |
| + | – | + | 0.013 (± 0.004) |
| + | + | + | 1.042 (± 0.016) |

**Table S5. CATNAP amplification loci of HPV-16.**

Sequence of CATNAP locus, highlighting the targeted PAM (pink) with central dipyrimidines (underlined) and protospacer sequence (red) and utilized by the indicated RNPs, intervening sequence length, and the shared Nt.BsmAI site (blue) used for amplification. Nt.BsmAI nicks the bottom strand, which serves as the target strand for the RNP.

| RNP | Sequence of locus |
|---|---|
| 3 | AGATACACCTACATTGCATGAATATAT---13nt--CCAGAGAC<br>TCTATGTGGATGTAACGTACTTATATA---13nt--GGTCTCTG |
| 5 | CAAGAACACGTAGAGAAACCCAGCTGT---52nt--CCAGAGAC<br>GTTCTTGTGCATCTCTTTGGGTCGACA---52nt--GGTCTCTG |
| 4 | GGGTCGGTGGACCGGTCGATGTATGTC---93nt--CCAGAGAC<br>CCCAGCCACCTGGCCAGCTACATACAG---93nt--GGTCTCTG |
| 6 | CAAAGACATCTGGACAAAAAGCAAAGA--134nt--CCAGAGAC<br>GTTTCTGTAGACCTGTTTTTCGTTTCT--134nt--GGTCTCTG |
| 7 | AGGTGTATTAACTGTCAAAAGCCACTG--176nt--CCAGAGAC<br>TCCACATAATTGACAGTTTTCGGTGAC--176nt--GGTCTCTG |
| 8 | TAGAACAGCAATACAACAAACCGTTGT--217nt--CCAGAGAC<br>ATCTTGTCGTTATGTTGTTTGGCAACA--217nt--GGTCTCTG |
| 9 | TGAGTATAGACATTATTGTTATAGTGT--258nt--CCAGAGAC<br>ACTCATATCTGTAATAACAATATCACA--258nt--GGTCTCTG |

**Table S6. HPV-type-specific CATNAP detection at 37°C.**

Mean (±SD) background-corrected *trans*-cleavage product (nM) from CATNAP reactions (**Figure 4A**) accumulated at 2.0 hours. Results are normalized and plotted in **Figure 4B**. Conditions: 37°C; 1.0 nM RNP, 0.5 pM HPV plasmid, amplification components, and 100 nM FQ-$C_{10}$ *trans*-substrate. Time-to-detect for the intended type defined as the time at which the signal (measured in the presence of amplification) exceeds a threshold corresponding to signal + 3SD (measured in the absence of amplification).

| RNP | HPV type-specificity | HPV type of target plasmid | | | | Time-to-detect (min) |
|---|---|---|---|---|---|---|
| | | 16 | 18 | 52 | 6b | |
| **4** | **16** | 76 (± 7) | -0.7 (± 0.2) | 1.2 (± 1.3) | 0.9 (± 0.6) | 0.3 |
| **84** | **18** | 0.5 (± 1.2) | 69 (± 4) | 0.8 (± 1.3) | 0.8 (± 0.7) | 4.3 |
| **85** | **52** | 2.4 (± 1.7) | 2.0 (± 1.9) | 84 (± 13) | 2.7 (± 0.8) | 2.3 |
| **86** | **6b** | -1.3 (± 1.0) | -0.1 (± 0.6) | 1.5 (± 1.7) | 70 (± 6) | 6.3 |

**Table S7. HPV-type-specific CATNAP detection at 25°C.**

Mean (± SD) background-corrected *trans*-cleavage product (nM) from CATNAP reactions (**Figure S11**) accumulated at 2.0 hours. Results are normalized and plotted in **Figure 4B**. Conditions: 25°C; 1.0 nM RNP, 0.5 pM HPV plasmid, amplification components, and 100 nM FQ-$C_{10}$ *trans*-substrate. Time-to-detect for the intended type defined as the time at which the signal (measured in the presence of amplification) exceeds a threshold corresponding to signal + 3SD (measured in the absence of amplification).

| RNP | HPV type-specificity | PAM | HPV type of target plasmid | | | | Time-to-detect (min) |
|---|---|---|---|---|---|---|---|
| | | | 16 | 18 | 52 | 6b | |
| **4** | **16** | AGAT | 33 (± 3) | 0.3 (± 0.2) | 0.1 (± 0.2) | 0.2 (± 0.1) | 0.3 |
| **84** | **18** | GAAT | 0.6 (±1.2) | 40(± 2) | -0.1 (± 0.1) | 0.1 (± 0.3) | 4.3 |
| **87** | **52** | GGGT | 0.3 (± 0.3) | 7.3 (± 0.1) | 43 (± 3) | 0.4 (± 0.2) | 2.3 |
| **86** | **6b** | AGGC | -0.1 (± 0.2) | 0.2 (± 0.2) | 0.6 (± 0.1) | 39 (± 1) | 6.3 |

**Table S8. Development of HPV type-specific RNPs for CATNAP.**

Apparent substrate turnover (values ± SE) obtained by fitting linear portion of background-corrected time courses (**Figure S10**) and normalizing slopes to target concentration. Conditions: 37°C; 1.0 nM RNP reacted with 2.5–5.0 pM target (either the plasmids carrying the HPV strains indicated or short 39–40-nt ssDNA) and 100 nM FQ-$C_{10}$ *trans*-substrate. RNPs were then used in type-specific CATNAP (**Figure 4**).

| RNP | HPV type-specificity | PAM | HPV plasmid | | Short ssDNA target |
|---|---|---|---|---|---|
| | | | Type | $k_{app}$ ($s^{-1}$) | $k_{app}$ ($s^{-1}$) |
| 4 | 16 | AGAT | 16 | -0.01 (± 0.01) | 4.45 (± 0.04) |
| 84 | 18 | GAAT | 18 | 0.11 (± 0.04) | 3.64 (± 0.05) |
| 85 | 52 | GAAC | 52 | 0.19 (± 0.05) | 2.72 (± 0.05) |
| 86 | 6b | AGGC | 6b | 0.02 (± 0.05) | 4.10 (± 0.06) |

**Table S9.** ***Trans*-substrate cleavage by AsCas12a RNP targeting HPV-16.**

Values (± SE) obtained by fitting the linear portion of time courses from **Figure S1**, expressed as background-corrected rates normalized to the target concentration approximation of activated RNP. Conditions: 37°C; 2.0 nM RNP reacted with 5.0 pM target (either the 858-bp pHPV or short 40-nt ssDNA) and 100 nM FQ-C10 *trans*-substrate. N/A = not applicable.

| RNP | PAM | ssDNA target $k_{app}$ ($s^{-1}$) | dsDNA target $k_{app}$ ($s^{-1}$) |
|---|---|---|---|
| None | - | | |
| 2 | ATTC | 2.88 (±0.02) | 1.14 (±0.03) |
| 3 | AGAT | 4.44 (±0.03) | -0.01(±0.01) |
| 10 | TGAT | 3.66 (±0.02) | 0.01 (±0.02) |
| 11 | CAGC | 1.61(±0.02) | 0.06 (±0.01) |
| 12 | AAAC | 1.61(±0.02) | 0.04 (±0.02) |
| 13 | AAGT | 2.99 (±0.04) | -0.04 (±0.02) |
| 14 | AACA | 2.24 (±0.02) | 0.06 (±0.01) |
| 15 | AAAC | 2.68 (±0.03) | -0.01 (±0.02) |
| 16 | GTGT | 5.08 (±0.04) | 0.02 (±0.01) |
| 17 | TAAA | 4.22 (±0.03) | 0.03 (±0.01) |
| 18 | CAGT | 3.44 (±0.04) | 0.03 (±0.01) |
| 19 | ATAC | 3.38 (±0.03) | 0.01 (±0.01) |
| 20 | GGCT | 2.90 (±0.02) | 0.04 (±0.02) |

**Table S10. Plasmids encoding Cas12a orthologs**

| Cas12a ortholog | Identifier |
|---|---|
| **Lb** | pIF162 |
| **As** | pIF468 |
| **Ts** | pIF1042 |
| **Fn** | pIF1038 |
| **Er** | pIF1039 |
| **Bo** | pIF1040 |
| **Bs** | pIF1041 |
